# Programmable CRISPRtune dissects the transcriptional repressive activity of MeCP2

**DOI:** 10.64898/2026.07.08.737375

**Authors:** Jinna I. Brim, Izaiah J. Ornelas, Peter J. Colias, Nikita S. Divekar, Da Xu, Justin P. Lubin, Sophia I. Ferrel, Luis Galán Palma, Mitzi G. Hernández Zamora, Rithu K. Pattali, Jameel J. McDaniel, Sarah E. Chasins, James K. Nuñez

## Abstract

The ability to control the expression of human genes is a major goal in synthetic biology, enables dissection of gene function, and can be harnessed for therapeutic applications. Advances in genome editing and transcriptional engineering often result in complete gene inactivation or full transcriptional repression. However, programmable tools to dial transcription at intermediate levels remain challenging. Here, we present CRISPRtune – a synthetic fusion of MeCP2 to catalytically dead dCas9 that tunes down transcription of endogenous genes in human cells by harnessing the mild repressor activity of MeCP2. Using pooled genome-scale CRISPR screens, we tune the expression of thousands of endogenous genes and define the targeting rules of CRISPRtune in human cells. With a platform to target MeCP2 at defined genomic sites, we show the direct epigenetic changes induced by MeCP2 at gene promoters and we identify its genetic dependency partners for productive transcriptional repression. Rett syndrome-associated mutations of MeCP2 show defects for transcriptional repression due to their failure to remodel the local epigenetic landscape of target genes. Together, we present a programmable method for transcriptional tuning in mammalian cells and offer an orthogonal platform to dissect the mechanistic function of chromatin regulators in living cells.

## INTRODUCTION

Transcription is regulated finely in mammals by several mechanisms, including transcription factors and proteins that participate in chromatin regulation. Thousands of genes in the human genome are dosage-sensitive, including around 3,000 estimated haploinsufficient genes^1^. Small reductions in gene expression can cause developmental disorders or disrupt cellular function, while others are more tolerant to changes in transcript levels^2,3^. For example, a variety of regulatory elements and genes are highly sensitive to SOX9 dosage, and misregulation of these targets is associated with craniofacial morphology defects and diseases like Pierre Robin sequence^4^. Furthermore, diseases that result from gain-of-function mutations, such as Huntington’s Disease, could be amenable to tuning to restore gene expression to healthy levels^5^. Thus, synthetic biology approaches to tune transcriptional output at precise levels advance our understanding of gene function, gene network mapping, disease modeling, and serve as potential therapeutic platforms^6^.

Genome editing platforms like CRISPR nucleases and prime/base editors induce genome modifications that lead to penetrant gene knockouts^7–9^. Transcriptional modifiers such as CRISPR interference (CRISPRi) rely on recruiting epigenome modifiers that induce strong transcriptional repression, thus limiting their use for applications that require more precise tuning of expression levels^10^. Recent efforts have discovered new effectors for programmable transcriptional repression; however, their generalizability and scalability across hundreds to thousands of gene targets in mammalian cells have not been characterized extensively^11–16^. Other platforms for programming tunable gene expression include RNA interference (RNAi), which is prone to off-target effects, and protein degradation tags, which currently is challenging to scale to hundreds or thousands of genes^17–20^. Additional strategies include incorporating microRNA-binding sites into an endogenous locus and mutating promoters and enhancers. These strategies require engineering at each target, which limits their scalability and ability to transfer across systems^21–23^. Recently, loading CRISPRi with attenuated single guide RNAs (sgRNAs) can program tunable gene expression at the transcriptional level; however, the platform can be heterogeneous across cells and the associated sgRNA mutations may introduce increased off-target effects^24^. Thus, a more generalizable, modular strategy is needed to achieve consistent intermediate repression across varying endogenous genes and cellular contexts.

Among the many proteins that mediate gene silencing in mammals is methyl-CpG binding protein 2 (MeCP2), a chromatin effector that binds DNA methylation via its methyl-CpG-binding domain (MBD) and recruits co-repressor complexes through its transcriptional repression domain (TRD). MeCP2 is most highly expressed in neurons, where it is present at levels approaching one molecule per nucleosome, an estimated 16 million molecules of MeCP2 per neuronal nucleus, highlights its importance in regulation transcription^25^. MeCP2 binds broadly throughout the genome at highly methylated regions of DNA and preferentially regulates expression of highly methylated long genes (>100 kb)^26–29^. MeCP2 represses transcription primarily by interacting with factors through its TRD that recruit histone deacetylases (HDACs) to remove acetyl groups from histone lysine residues^30,31^. MeCP2 interacts with the HDAC-SIN3A co-repressor complex, as well as the nuclear receptor co-repressor (NCoR)-silencing mediator for retinoic acid and thyroid hormone receptor (SMRT) complex, which contains HDAC3^32^. Mutations in the *MECP2* gene are associated with Rett syndrome (RTT), a neurodevelopmental disorder linked to more than 900 pathogenic variants^33–35^. Many RTT-associated mutations cluster primarily in the methyl-binding domain (MBD) and transcriptional-repression domain (TRD), yet the precise mechanism of how distinct mutations affect disease phenotypes remains poorly understood^36,37^.

Previously, MeCP2 has been harnessed for CRISPR-based programmable transcriptional repression, typically as a synthetic fusion with a KRAB domain (KRAB-MeCP2)^12,38–42^. The combinatorial repressor can induce potent gene silencing in cell lines and mouse models at levels higher than CRISPRi-KRAB. Although fusions of MeCP2 to dCas9 have been tested on synthetic gene reporters and a limited number of endogenous genes, the programmable repressive activity of MeCP2 remains unclear and its generalizability across endogenous human genes has yet to be profiled^11,12,43^.

Here, we present a synthetic fusion of MeCP2 to dCas9 called CRISPRtune, a platform for tuning gene expression to intermediate levels by harnessing the repressive activity of MeCP2. We show that CRISPRtune is functional across different human cell types and define the epigenetic changes induced by CRISPRtune at target loci. Further, we show that incorporating Rett syndrome-associated mutations into CRISPRtune leads to defective transcriptional repression. Finally, we utilize pooled CRISPR screens to tune the expression of thousands of essential genes and empirically define sgRNA targeting rules for effective tuning, demonstrating its broad applicability and tunability across essential genes. Together, our findings establish CRISPRtune as a tool for regulating gene dosage and as a platform to dissect the molecular function of chromatin regulators at defined loci in living cells.

## RESULTS

### MeCP2-dCas9 tunes gene expression

Previously, the C-terminal region of the rat MeCP2 (rMeCP2) was fused to dCas9 for programmable transcriptional repression of a synthetic reporter in mammalian cells^44^. We first constructed a single fusion construct of the C-terminal region of the rMeCP2 to the N-terminus of catalytically dead Cas9 (dCas9) (**Fig. 1a**). Human and rat MeCP2 share 94% protein sequence similarity (**Supplementary Fig. 1a**). The C-terminal region contains the transcriptional repressor domain (TRD) and excludes the MBD domain, which allows us to decouple the repression activity of MeCP2 from its DNA binding MBD domain.

**Figure 1.**
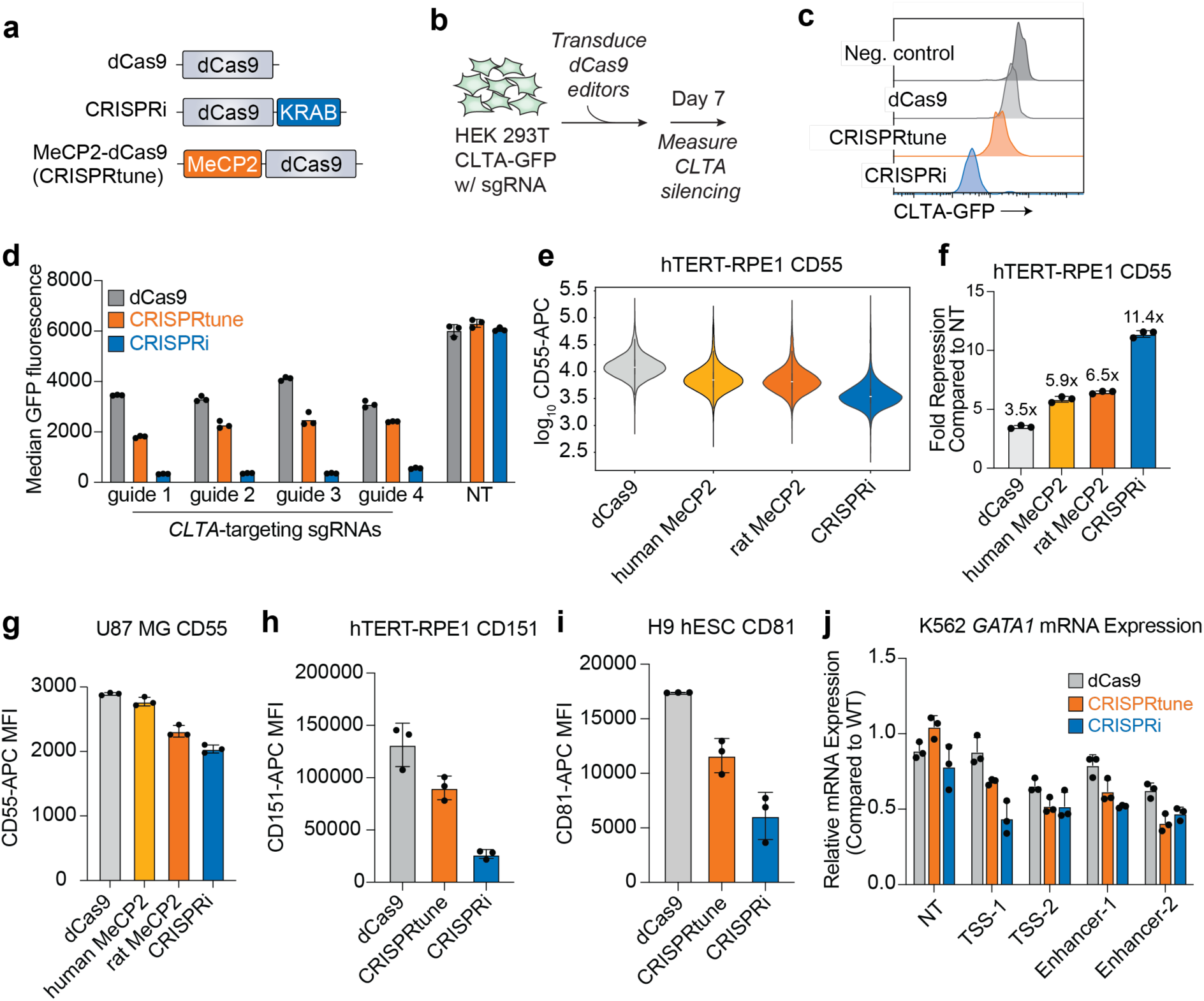
Programmable tuning of endogenous genes with CRISPRtune. **(a)** A schematic of the dCas9 constructs used in this study. **(b)** Experimental workflow to test the repressive activity of dCas9, CRISPRi, and CRISPRtune by generating HEK293T cell lines, each expressing a dCas9 repressor targeting the *CLTA* gene, followed by quantitative CLTA-GFP protein measurement in single cells by flow cytometry. **(c)** A histogram plot comparing CLTA protein levels in cells expressing dCas9, CRISPRi, or CRISPRtune targeting the endogenously GFP-tagged *CLTA* gene. **(d)** Quantification of median GFP fluorescence in CLTA-GFP cells across four sgRNAs targeting *CLTA* and a non-targeting control sgRNA. **(e)** A violin plot comparing *CD55* expression levels in hTERT-RPE1 single cells measured by cytometry with dCas9 editor fusions. The data are aggregated from three technical replicates. **(f)** Quantification of *CD55* expression across editors in hTERT-RPE1 cells with an sgRNA targeting *CD55* normalized to a non-targeting sgRNA for each editor. Data represent the mean ± s.d. (*n*=3 technical replicates). Human MeCP2, rat MeCP2, and CRISPRi repress *CD55* to a stronger degree than dCas9 alone (*p* < .05). **(g)** Quantification of *CD55* expression in U-87 MG cells. *CD55* repression differs from dCas9 with rat MeCP2 (*p* = .0003) and is not significantly different from dCas9 with human MeCP2 (*p* = .2149). **(h)** Quantification of *CD151* expression in hTERT-RPE1 cells. *CD151* repression by CRISPRtune is significantly different than by dCas9 (*p* = .0003) and CRISPRi (*p* = .0456). **(i)** Quantification of *CD81* expression in H9 human embryonic stem cells. *CD81* protein levels are significantly different when *CD81* is targeted with CRISPRtune compared to dCas9 (*p* = .0031) and CRISPRi (*p =* .0229). **(j)** Relative mRNA expression of *GATA1* measured by RT-qPCR with CRISPRtune (orange) or CRISPRi (blue) and five different sgRNAs: non-targeting (NT), two different sgRNAs targeting the transcription start site (TSS), and two different sgRNAs targeting the enhancer.

To test its programmable repression activity, we transfected plasmids encoding MeCP2-dCas9 into HEK293T cells expressing the endogenous *CLTA* gene tagged with GFP^45,46^. The cells constitutively express three distinct sgRNAs that tile across the *CLTA* promoter (**Supplementary Fig. 1c**). In parallel, we compared MeCP2-dCas9 to dCas9 and CRISPRi (KOX1-KRAB)^47^. Two days after transient transfection, we sorted for cells that express the dCas9 fusions and monitored CLTA-GFP protein levels with flow cytometry as a readout of transcriptional repression activity. At four days post-transfection, each construct represses CLTA-GFP in over 90% of sorted cells (**Supplementary Fig. 1d**). However, by eight days post-transfection, the degree of CLTA protein repression (measured by median CLTA-GFP intensity) weakens drastically for dCas9, remains strong for CRISPRi, and exhibits intermediate repression for MeCP2 relative to dCas9 and CRISPRi **(Supplementary Fig. 1e, f).** CLTA protein levels revert to pre-edited expression levels by 15 days as the plasmids expressing the dCas9 fusions are no longer present in cells, indicating that these repressors do not induce long-term epigenetic silencing (**Supplementary Fig. 1d**).

Transient plasmid transfections result in varying degrees of dCas9-repressor expression across the transfected cell population that can introduce heterogeneity across the cell population. Thus, we next introduced the dCas9 repressors by lentiviral transductions for constitutive expression. In sgRNA-expressing CLTA-GFP lines, we transduced HEK293T cells with lentiviral particles encoding MeCP2-dCas9, CRISPRi, or dCas9 (**Fig. 1b**). Seven days post-transduction, CRISPRi represses CLTA strongly, whereas MeCP2-Cas9 lowers CLTA expression at levels intermediate between dCas9 and CRISPRi (**Fig. 1c, d**). We further tested a total of four different sgRNAs that bind to the promoter of *CLTA* and a non-targeting sgRNA negative control. We detect intermediate CLTA levels across all four sgRNAs with MeCP2-dCas9, whereas CRISPRi consistently represses CLTA strongly. Similarly, when delivered as mRNA, MeCP2-dCas9 results in repression intermediate between dCas9 and CRISPRi (**Supplementary Fig. 1g**). Given these observations, we hereafter refer to our MeCP2-dCas9 construct as CRISPRtune.

### CRISPRtune is effective across different genes and cell types

To determine the generality of CRISPRtune activity across different cell types, we tested CRISPRtune in human retinal pigment epithelial (hTERT-RPE1) cells. We generated cell lines that constitutively express CRISPRi, CRISPRtune, and dCas9. We then introduced an sgRNA through lentiviral transduction targeted to the promoter of *CD55,* a nonessential gene for cell viability, and measured cell surface-localized CD55 protein levels using flow cytometry. CRISPRtune induces a 6.5-fold repression in CD55 levels compared to a non-targeting sgRNA, a level intermediate to the 3.5-fold repression by dCas9 (*p* < .0001) and 11.4-fold repression by CRISPRi (*p* < .0001) (**Fig. 1e, f**). Next, we compared the repressive activity of the human and rat orthologs of MeCP2. The human MeCP2 fusion to dCas9 results in a 5.9-fold repression of *CD55*, whereas the original rat MeCP2 fusion results in a 6.5-fold repression (**Fig. 1f**). These data suggest the rat and human orthologs of MeCP2 repress gene expression at similar levels, likely due to their high sequence similarity (**Supplementary Fig. 1a**).

MeCP2 is expressed in all human cell types but is most abundantly expressed in neuronal and glial cells where it regulates neuron-specific gene expression^37^. We thus tested the repressive activity of CRISPRtune in U87-MG glioblastoma cells and observed that CRISPRtune lowers CD55 expression at levels intermediate between dCas9 (*p* = .0033) and CRISPRi (*p* = .0003) (**Fig. 1g**). We further tested CRISPRtune silencing on additional cell surface genes *CD151*, *CD81*, and *CD29* in hTERT-RPE1 cells, and similarly observed tuning of gene expression (**Fig. 1h, Supplementary Fig. 2a-b**). CRISPRtune similarly tunes gene expression in H9 human embryonic stem cells when targeted to *CD81* and *CD151* (**Fig. 1i, Supplementary Fig. 2c**). These observations support that the transcriptional tuning activity of CRISPRtune can be harnessed across genes and cell types.

To further distinguish the performance of CRISPRtune from baseline dCas9 steric hinderance, we constructed a truncation of MeCP2 fused to dCas9 and compared its performance to CRISPRtune in hTERT-RPE1 cells targeting *CD55*^38^. The truncation comprises of an 80 amino acid region of the transcriptional repression domain (**Supplementary Fig. 2d**). We observe similar levels of *CD55* repression with truncated MeCP2 as with the CRISPRtune construct that comprises a 298 amino acid region of MeCP2 (**Supplementary Fig. 2e)**. Together, these data differentiate the performance of CRISPRtune from existing tools and demonstrate its applicability across genes and cell types.

### CRISPRtune programs transcriptional tuning of enhancers

Next, we evaluated whether CRISPRtune could modulate gene expression when directed to distal regulatory elements. We targeted a well-characterized *GATA1* enhancer using previously validated sgRNAs for the transcriptional start sites (TSS-1 and TSS-2), enhancer sites (Enhancer-1 and Enhancer-2), and a non-targeting control in K562 cells^48,49^. *GATA1* expression was measured in cells expressing CRISPRtune or CRISPRi by reverse transcription quantitative polymerase chain reaction (RT-qPCR). As expected, cells expressing CRISPRi induces stronger repression when targeted to the TSS and enhancer, with TSS-1 and TSS-2 reducing GATA1 expression by 56% and 48%, respectively, and Enhancer-1 and Enhancer-2 reducing expression by 48% and 53%, respectively (**Fig. 1j**). Meanwhile, CRISPRtune programs more moderate decrease in expression, with the TSS-1 and TSS-2 reducing GATA1 levels by 32% and 48%%, respectively, and Enhacer-1 by 38%. Surprisingly, CRISPRtune targeting to Enhancer-2 resulted in stronger repression than CRISPRi, reducing GATA1 expression by 59%, suggesting that transcriptional outcomes may depend on sgRNA position of the epigenome editor. These results indicate that CRISPRtune can modulate gene expression by targeting enhancer elements, expanding tunable transcription control to distal regulatory elements.

### Transcriptional and epigenetic changes induced by CRISPRtune

To examine how CRISPRtune modulates transcription at the chromatin level, we profiled transcriptomic and epigenetic changes at target gene promoters. We performed RNA-sequencing (RNA-seq) in hTERT RPE-1 cells expressing dCas9, CRISPRi, and CRISPRtune targeting CD55 and a non-targeting sgRNA control. RNA-seq analysis revealed that CD55 transcript abundance was reduced by 2.39-fold for dCas9, while CRISPRtune induced a 3.87-fold repression and 31.69-fold repression for CRISPRi (**Fig. 2a, b**). CRISPRtune also shows high specificity, with one off-target transcript showing a log2 fold-change >1 and adjusted p-value <0.5 (**Supplementary Fig. 3a-c**).

**Figure 2.**
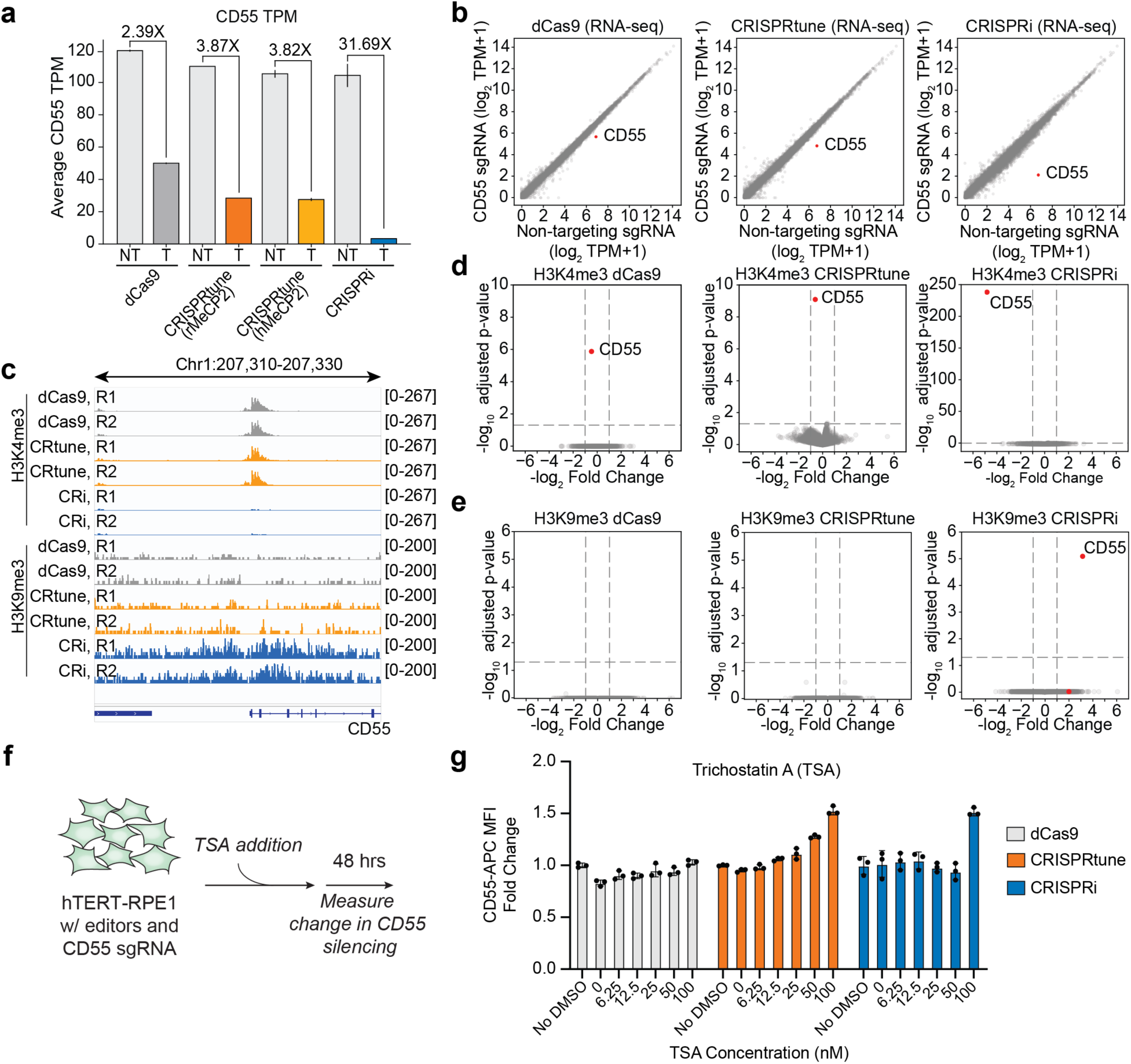
Transcriptional tuning and epigenomic changes induced by CRISPRtune. **(a-e)** RNA-seq and CUT&RUN plots of hTERT-RPE1 cells following lentiviral transduction of dCas9, CRISPRtune, and CRISPRi, then lentiviral transduction of *CD55* and NT sgRNAs. **(a)** Quantification of CD55 transcripts per million (TPM) from RNA-seq in hTERT-RPE1 cells. Data represent the mean of *n*=2 technical replicates. **(b)** A comparison of *CD55* TPM between a non-targeting sgRNA (x axis) and CD55 sgRNA (y axis) for dCas9 (left), CRISPRtune (middle), and CRISPRi (right) from RNA-seq. Data represent average TPM for *n*=2 technical replicates. **(c)** A view of a 20 kb genomic window containing the *CD55* locus and CUT&RUN sequencing results for H3K4me3 and H3K9me3. Tracks for two technical replicates are labeled R1 and R2 for dCas9 (gray), CRISPRtune (orange), and CRISPRi (blue). **(d,e)** Volcano plots comparing H3K4me3 (**d**) and H3K9me3 (**e**) CUT&RUN sequencing data between dCas9, CRISPRtune, and CRISPRi with either *CD55*-targeting or NT sgRNAs. Red dots represent *CD55*. No peaks were called for H3K9me3 in the dCas9 and CRISPRtune samples. **(f)** A schematic of the experimental workflow for addition of a pan-HDAC inhibitor, Trichostatin A (TSA). TSA was added in a range of doses, resuspend and diluted in the same volume of DMSO to hTERT-RPE1 cells actively silencing *CD55*. **(g)** Quantification of *CD55* expression in hTERT-RPE1 cells. Data are represented as fold change normalized to CD55 protein levels in the no DMSO control for each respective editor.

Next, we profiled whether the observed transcriptional changes are associated with differences in local chromatin environments. We performed CUT&RUN to profile changes to transcriptionally repressive H3K9me3 and activating H3K4me3 marks in CRISPRi, CRISPRtune, and dCas9 expressing hTERT-RPE1 cells targeting *CD55* (**Fig. 2c-e**). Cells expressing dCas9 alone minimally reduces H3K4me3 and do not enrich for H3K9me3. Meanwhile, CRISPRi greatly depletes H3K4me3 and enriches for H3K9me3, consistent with its strong silencing activity. In contrast, CRISPRtune shows modest but increased reduction in H3K4me3 compared to dCas9 without enrichment for H3K9me3. These data suggest that CRISPRtune and CRISPRi function through two different mechanisms that result in their respective tuning and repressing activities.

To explore epigenetic changes induced by MeCP2, we investigated the role of histone deacetylases (HDACs) in CRISPRtune mediated silencing. MeCP2 primarily represses transcription by interacting with factors through its transcriptional repression domain that recruit HDACs to remove acetyl groups from histone lysine residues^31,50^. We thus examined the effect of a pan-HDAC inhibitor, Trichostatin A (TSA), on CRISPRtune based repression of *CD55* (**Fig. 2f**). We observed a dose-dependent loss in *CD55* repression by CRISPRtune 48 hours after TSA addition, suggesting TSA inhibits the role of HDACs in CRISPRtune mediated silencing (**Fig. 2g)**. TSA did not inhibit repression by dCas9 alone, supporting the mechanism that MeCP2 recruits HDACs to remove histone acetyl marks to silence gene expression. TSA addition did not inhibit CRISPRi based repression at low doses; however, we observe a loss in repression at the highest dose of 100 nM. This observation is consistent with the known role of KRAB to recruit TRIM28, which serves as a scaffold for the NuRD complex that contains HDACs^51^. These results suggest that CRISPRtune mediated silencing is dependent on the recruitment of HDACs to tune gene expression.

### Rett syndrome-associated MeCP2 mutants are defective in repression

The transcriptional repression domain (TRD) of MeCP2 is a well-known hotspot for mutations that correlate with a wide range of symptoms and severity in Rett syndrome patients^33,52^. RTT mutations in the TRD have been shown to disrupt the ability of MeCP2 to interact with co-repressors SIN3A and NCoR/SMRT and impair the ability of MeCP2 to repress transcription^52,53^. Engineered dCas9-based epigenome editors have been used as a platform to examine the functional effects of mutations in chromatin modifying enzymes^54^. To test the effect of Rett syndrome-associated mutations on CRISPRtune-mediated repression, we engineered RTT missense mutations in the MeCP2-TRD region of CRISPRtune (**Fig. 3a**). We selected variants with high clinical frequency from RettBASE, an MeCP2 variant database for RTT and related clinical phenotypes^55^. We selected variants with a range of AlphaMissense pathogenicity scores, where a score closer to 1 indicates a higher probability of being pathogenic and a score closer to 0 indicates a benign variant: K304E (0.998), R306C (0.997), P225R (0.97), P302R (0.993), K305R (0.808), V300I (0.23), A279V (0.092).

**Figure 3.**
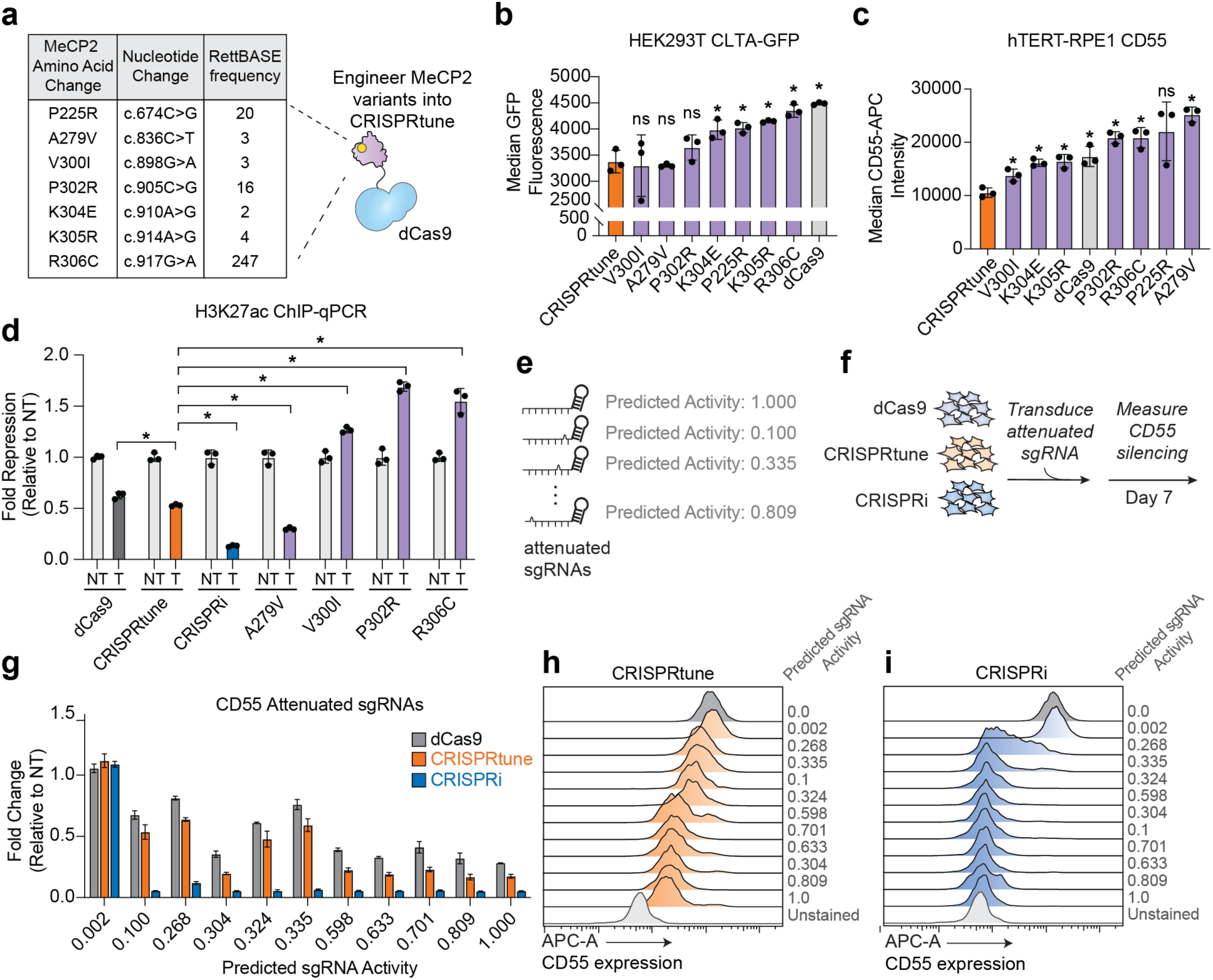
Rett-Syndrome associated MeCP2 mutants and attenuated sgRNAs titrate CRISPRtune mediated repression. **(a)** A summary of MeCP2 variants engineered into CRISPRtune, including the amino acid change, nucleotide change, and RettBASE frequency. RettBASE frequency is the number of times a mutation was identified and listed in the repository. **(b)** Quantification of median GFP fluorescence of HEK293T cells expressing CRISPRtune and CRISPRtune engineered with common RTT variants. Cells constitutively express an sgRNA targeting *CLTA*. Significant p-values (*p* < .05) from a two-sided unpaired t-test with CRISPRtune are denoted with (*). Non-significant p-values (*p* > .05) are denoted with (ns). **(c)** Quantification of median CD55-APC fluorescence of hTERT-RPE1 cells expressing CRISPRtune and CRISPRtune engineered with common RTT variants. Cells constitutively express an sgRNA targeting *CD55*. **(d)** Quantification of H3K27ac fold enrichment from ChIP-qPCR at *CD55*. H3K27ac fold enrichment for an editor targeting CD55 was normalized to a non-targeting sgRNA. Significant p-values (*p* < .05) from a two-sided unpaired t-test with CRISPRtune are denoted with (*). **(e)** A schematic of attenuated sgRNA design. A predicted activity of 1.000 is an sgRNA without modifications. **(f)** Experimental workflow for testing attenuated sgRNAs with CRISPRtune in hTERT-RPE1 cells. sgRNAs were selected with a range of predicted activities. **(g)** Quantification of fold repression of CD55 with a range of attenuated sgRNAs relative to a non-targeting sgRNA. **(h,i)** A histogram plot of flow cytometry data from hTERT-RPE1 cells that stably express CRISPRtune (**g**) or CRISPRi (**h**) and an attenuated sgRNA.

CRISPRtune variants were introduced by lentiviral transduction in sgRNA-expressing CLTA-GFP HEK293T lines (**Fig. 3b**). CRISPRtune reduces expression 2.4-fold, while CRISPRtune-K304E, P225R, K305R, and R306C resulted in 2.03, 2.01, 1.95, 1.86-fold repression, respectively. Notably, R306C is the most common RTT mutation in the TRD recorded in patients^33,34,57^, and CRISPRtune-R306C is the most defective variant in repressive activity with a 22.5% loss in repression compared to CRISPRtune at levels similar to dCas9 only.

We further tested Rett syndrome-associated variants in hTERT-RPE1 cells to determine if mutations disrupt CRISPRtune-mediated repression in a different cell type. Similarly, all tested variants tested resulted in loss of CD55 silencing (**Fig. 3c**). Notably, variants P302R, R306C, P225R, and A279V resulted in weaker silencing than dCas9 in hTERT-RPE1 cells. Common Rett variants R306C and P225R performed similarly across cell types and genes, with both resulting in the highest loss in CRISPRtune-based repression of CLTA-GFP in HEK293T and CD55 in hTERT-RPE1 cell lines, which may be correlated to their high clinical severity score.

Mutations in MeCP2 have been shown to disrupt histone deacetylation of H3K27^26^. We thus sought to profile changes to H3K27ac of cell lines with RTT variants engineered in CRISPRtune. We performed chromatin immunoprecipitation followed by quantitative polymerase chain reaction (ChIP-qPCR) on hTERT-RPE1 cells constitutively expressing CRISPRtune variants K304E, P225R, K305R, and R306C, as well as WT CRISPRtune, CRISPRi, and dCas9. CRISPRi lines targeting *CD55* induce a 7.6-fold decrease in H3K27ac levels relative to a non-targeting sgRNA (**Fig. 3d**). CRISPRtune induces a 1.9-fold decrease in H3K27ac levels, while dCas9 alone results in a 1.6-fold decrease. Variants V300I, P302R, and R306C engineered in CRISPRtune targeting *CD55* results in higher levels of H3K27ac than WT CRISPRtune, with H3K27ac levels higher than a non-targeting sgRNA for each editor. This data corroborates previous reports that the common RTT variant R306C is associated with an increase in H3K27ac levels of MeCP2-bound genomic loci, though the mechanism remains unclear^25,26^.

Altogether, these data support that RTT mutations disrupt transcriptional repression by MeCP2. Furthermore, these data highlight that introducing variants into CRISPRtune further expands the dynamic range of gene expression tuning. More broadly, our set of CRISPRtune variants introduces a new set of fusion proteins that can be used to fine tune gene expression to investigate gene dosage sensitivities.

### Attenuated sgRNAs expand dynamic range of tuning with CRISPRtune

We next sought additional strategies to further broaden the dynamic range of transcriptional tuning with CRISPRtune. We employed a previously described approach that couples CRISPRi and attenuated sgRNAs^24^. This strategy rationally introduces mutations within the sgRNA protospacer sequence, ultimately resulting in graded transcriptional repression of endogenous genes by CRISPRi. We benchmarked CRISPRtune with CRISPRi loaded with attenuated sgRNAs by selecting 12 sgRNAs targeting *CD55* with predicted activities between 0 and 1.0, where 0 is a non-targeting control and 1.0 is the sgRNA with the best predicted repression activity for CRISPRi (**Fig. 3e**). We then introduced sgRNAs through lentiviral transduction into hTERT-RPE1 cells constitutively expressing CRISPRtune and CRISPRi (**Fig. 3f**).

Notably, across all attenuated sgRNAs tested, CRISPRtune consistently represses gene expression at levels between dCas9 and CRISPRi (**Fig. 3g**). Nearly all sgRNAs tested with CRISPRi silence CD55 expression at similar levels as an sgRNA with the strongest predicted repressive activity. An exception is an sgRNA with predicted activity of 0.268 that results in heterogeneous silencing across the cell population (**Fig. 3i**). In contrast, we observe more graded repression with CRISPRtune, with the predicted optimal sgRNA resulting in the strongest repression and the predicted least optimal sgRNA producing the weakest repression, similar to a non-targeting sgRNA control (**Fig. 3h**). Ranking attenuated sgRNA activities for CRISPRtune reveals a graded tuning of CD55 levels that further illustrates the ability to attenuate CRISPRtune repression. Moreover, the use of attenuated sgRNAs with dCas9 alone also leads to graded tuning of CD55 levels, although the range of suppression is more limited than with CRISPRtune (**Supplementary Fig. 4a**). Collectively, these results demonstrate that CRISPRtune provides a platform for intermediate levels of repression and can be integrated with existing sgRNA design strategies to enable more refined control of transcriptional output.

### Targeting principles for MeCP2-mediated transcriptional repression

Defining the optimal sgRNA for CRISPR-based technologies is critical for their effective use. For CRISPR transcriptional repressors and activators, functional sgRNAs can be derived empirically by tiling sgRNAs across target gene promoters and identifying sgRNAs that are depleted or enriched using phenotypic readouts such as cell viability^10,58^. Previously, sgRNA tiling screens were performed with dCas9-KRAB-MeCP2 that yielded the same optimal sgRNA targeting rules as dCas9-KRAB^12^. Since both platforms utilized KRAB, it is unclear whether MeCP2 alone has a similar targeting profile as KRAB.

Using sgRNA tiling screens, we sought to determine the preferential binding site of MeCP2-dCas9 for repressive activity across endogenous gene promoters. Although cell growth is nonlinear, targeting essential genes can be used as a proxy for repressor strength and probes the generality of the platform^59^. We curated a list of 1,869 genes that are essential for cancer cell viability from the Cancer Dependency Map (DepMap)^60^ and designed sgRNAs that tile across a +/− 1–2.5 kilobase (kb) region relative to the transcriptional start site (TSS), totaling 472,633 unique sgRNAs. We calculated phenotype (γ) scores by comparing sgRNA abundance at T_final_ to T_initial_ after 11 population doublings in K562 cells expressing CRISPRtune^61^ (**Fig. 4a**). We performed two technical replicates of the sgRNA tiling screens that resulted in high correlation in γ scores for each sgRNA across the two replicates (**Supplementary Fig. 5a, b**). Violin plots comparing phenotype scores for targeting against non-targeting sgRNAs show growth defects for essential genes, confirming that the tiling screen effectively identifies sgRNAs with strong growth phenotypes (**Supplementary Fig. 5c, d**).

**Figure 4.**
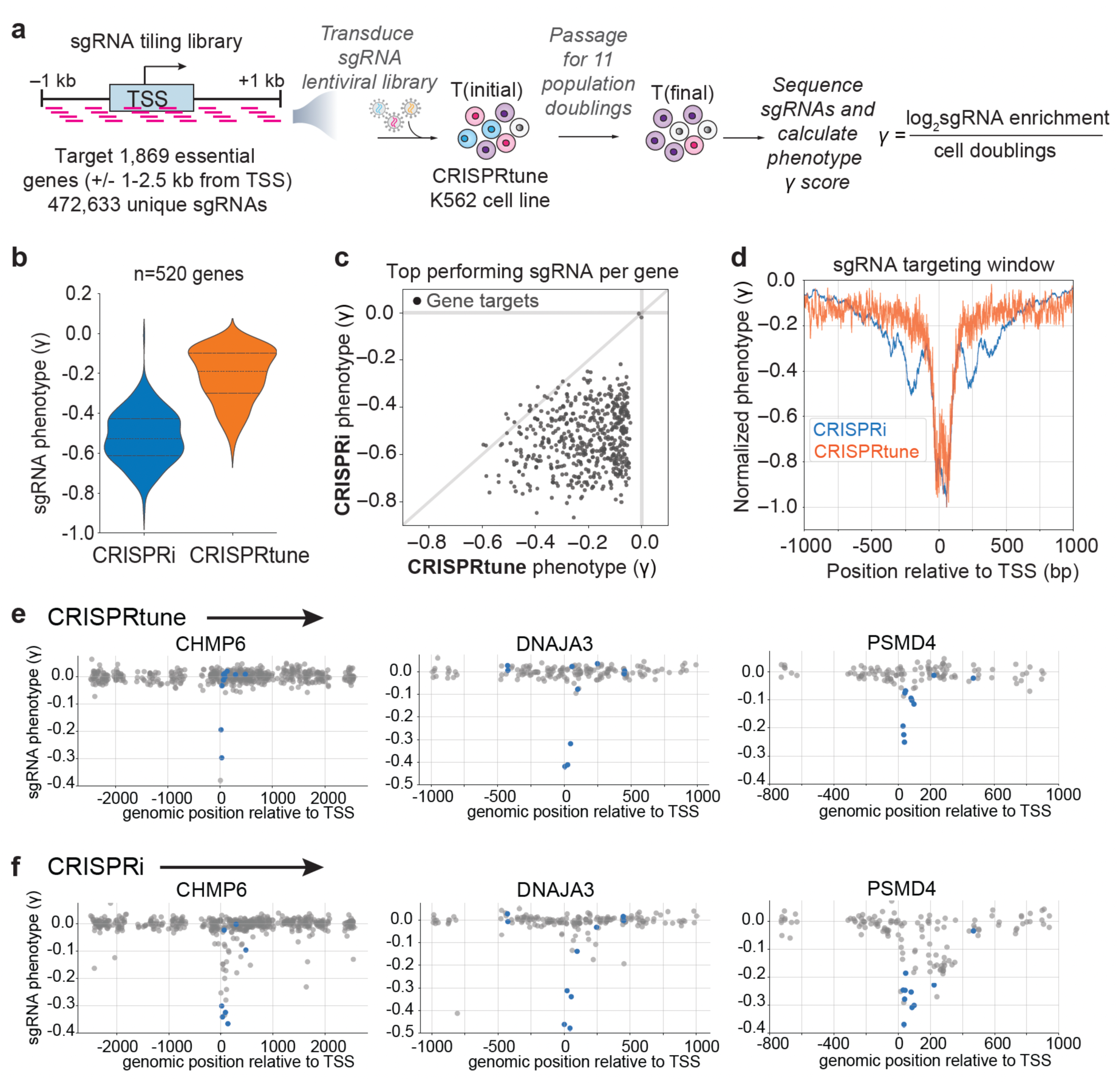
Pooled sgRNA tiling screen defines optimal sgRNA rules for CRISPRtune. **(a)** A schematic of the sgRNA tiling screen workflow. The sgRNA library tiles a region ±1–2.5 kb from the transcription start site (TSS). **(b)** A comparison of the phenotype score (γ) for all sgRNAs between CRISPRi and CRISPRtune. **(c)** A comparison of the phenotype score (γ) for the top performing sgRNA per gene between CRISPRtune (x axis) and CRISPRi (y axis). Each dot represents one sgRNA. **(d)** An aggregate plot comparing the normalized phenotype score for each sgRNA. The orange line represents screen data from CRISPRtune and the blue line represents screen data from CRISPRi. **(e,f)** Representative plots of screen data from CRISPRtune (**e**) and CRISPRi (**f**) at three genes: *CHMP6*, *DNAJA3*, and *PSMD4*. Each dot represents one sgRNA tested in the screen and blue dots represent the best computationally predicted sgRNA for CRISPRi.

Previously, we performed a tiling sgRNA screen for 520 essential genes with CRISPRi (**Supplementary Table 4**)^46^. Using this gene set, we compared the growth phenotype and optimal sgRNA performance between CRISPRi and CRISPRtune. For initial analysis, we selected the top-performing sgRNA per gene as the preferred top sgRNA per editor may differ. We observe similar trends in phenotype strength between CRISPRi and CRISPRtune, where CRISPRi induces stronger growth defects, while CRISPRtune yields more moderate growth defects (**Fig. 4b, c**). To derive optimal sgRNAs for CRISPRtune, we normalized the phenotypes across the screens and generated an aggregate plot of sgRNA activities relative to the TSS (**Fig. 4d**). CRISPRi displays a broader range of activity extending upstream and downstream of the TSS, as reported previously^61^. In contrast, functional CRISPRtune sgRNAs are more narrowly restricted around the TSS (**Fig. 4d**). These findings suggest that MeCP2-dCas9-mediated repression is more position-dependent than CRISPRi, with repression tightly centered around the TSS. In contrast, CRISPRi exhibits a broader window of activity that extends both upstream and downstream, as demonstrated in representative gene-level profiles for *CHMP6*, *DNAJA3*, and *PSMD4* (**Fig. 4e, f**).

Notably, the best-performing sgRNAs for CRISPRtune are also within the top ten predicted optimal sgRNAs for CRISPRi and induce stronger growth defects with CRISPRi than with MeCP2 (**Supplementary Fig. 5e**). These results indicate that CRISPRi-optimized sgRNAs are also effective with MeCP2, highlighting the utility of existing CRISPRi sgRNA libraries for CRISPRtune-mediated gene repression. More broadly, our screen is the largest CRISPR sgRNA tiling screen of protein-coding genes to our knowledge, and our tiling sgRNA dataset provides users with pre-validated sgRNAs for productive transcriptional tuning for 1,869 endogenous essential genes.

### Generality of CRISPRtune across essential genes

As demonstrated previously with CRISPRi loaded with attenuated sgRNAs, tuning the expression of essential genes at intermediate levels can result in measurable defects in cell proliferation over time^24^. Thus, we evaluated the generality of CRISPRtune by performing pooled CRISPR screens to target genes that are essential for cell viability, using cell depletion as a proxy for transcriptional repression strength^13,48,62,63^. We compiled a set of 1,837 genes classified as essential for cancer cell viability based on the Cancer Dependency Map (DepMap)^60,64^. For each essential gene, we selected four sgRNAs from previously published CRISPRi libraries^39,58^. We included sgRNAs targeting non-essential genes and non-targeting controls, resulting in a final library of 8,086 sgRNAs that we term the Ornelas essential gene library (**Fig. 5a**)^65^. We transduced the Ornelas library into K562 cells expressing dCas9, CRISPRtune, or CRISPRi, passaged the cells for ∼11 population doublings, and calculated a γ score by comparing sgRNA abundance at the final time point (T_final_) to the initial population (T_initial_) (**Fig. 5b and Supplementary Fig. 7a-f**).

**Figure 5.**
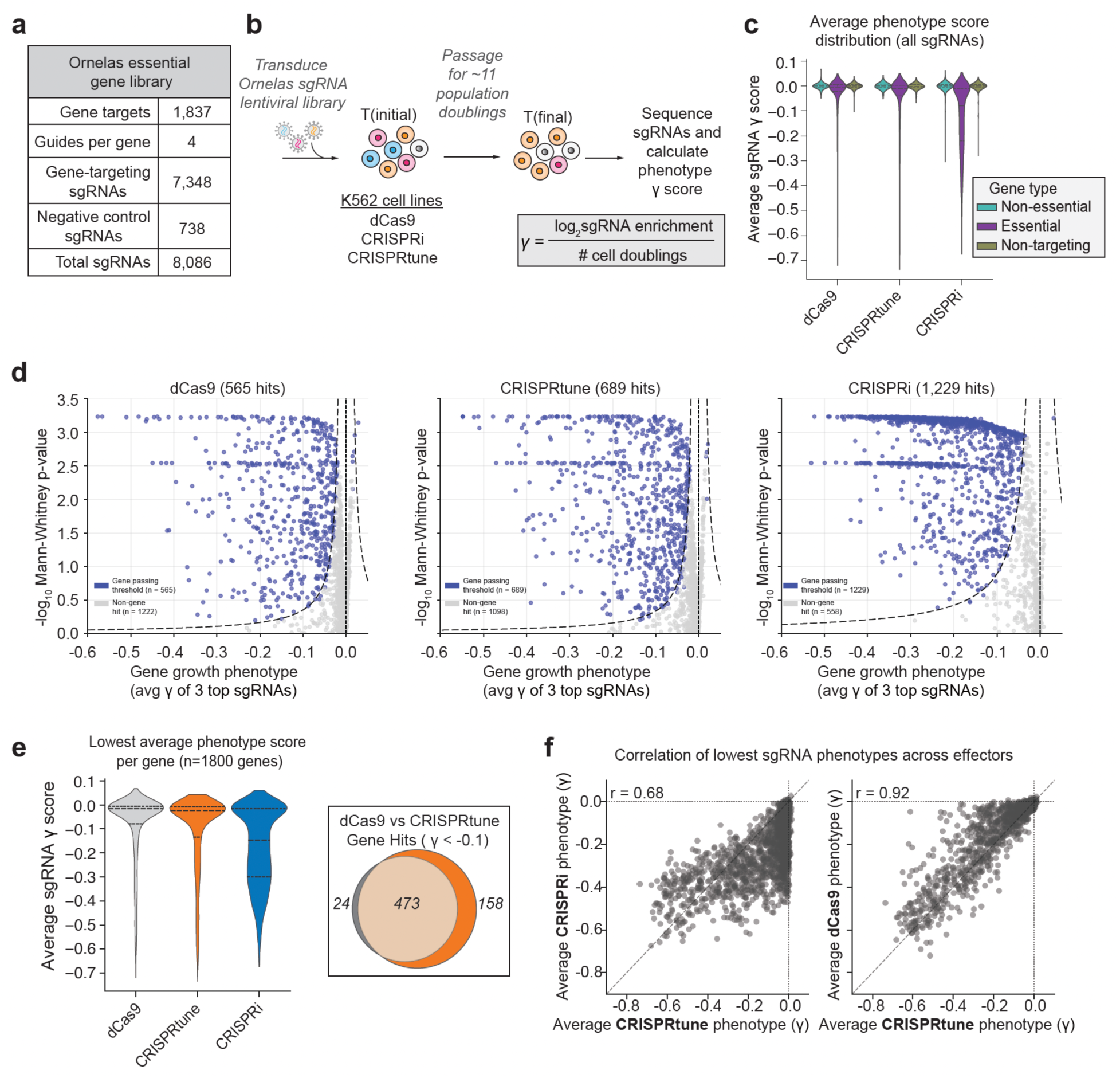
CRISPRtune modulates essential gene phenotypes. **(a)** A summary of the number of genes and sgRNAs per indicated category in the essential gene library. **(b)** A schematic of the essential gene library screen workflow. **(c)** A violin plot of the average sgRNA phenotype score (γ) for dCas9, CRISPRtune, and CRISPRi. Populations are grouped by sgRNAs for nonessential genes (teal), essential genes (purple), and non-targeting (yellow). **(d)** Volcano plots comparing the average gene growth phenotype of 3 top sgRNAs (x axis) and the -log10 Mann-Whitney p-value. Blue dots represent genes passing the threshold of significance and gray dots represent non-gene hits. **(e)** A violin plot of the lowest average phenotype scores for each gene (left). A Venn diagram comparing the number of gene hits (γ < 0.1) for dCas9 (gray) and CRISPRtune (orange). **(f)** A comparison of the lowest sgRNA phenotype for each sgRNA between CRISPRtune and both CRISPRi (left) and dCas9 (right).

We first compared the distribution of γ scores across all the three CRISPR editors. Across essential genes, CRISPRi induced the strongest growth defects, followed by CRISPRtune and dCas9, respectively (**Fig. 5c**). To assess gene-level effects across effectors, we computed a Mann-Whitney p-value comparing the γ scores of all sgRNAs targeting essential genes to those of the non-targeting control sgRNAs as previously reported. Additionally, we calculated a z-score for each gene based on the average phenotype of its top three sgRNAs relative to the distribution of non-targeting controls. Using an empirically derived threshold combining z-score and p-value, we quantified the number of gene-level hits for each effector. This analysis reveals distinct repression strengths for each effector, with CRISPRi producing the strongest growth phenotypes (1,229 hits), followed by CRISPRtune (689 hits) and dCas9 (565 hits) (**Fig. 5d**).

Furthermore, we selected the top-performing sgRNA per gene independent of effector and observe that CRISPRtune phenotypes are stronger than those of dCas9, as highlighted by a downward shift in γ scores, but weaker than CRISPRi, suggesting a tunable mode of repression (**Fig. 5e**). Importantly, CRISPRtune represses more genes than dCas9, with ∼25% more genes showing growth defects (γ < –0.1). Finally, we assessed the correlation of phenotypes across effectors by correlating the average phenotype of the most depleted sgRNA per gene.

CRISPRtune and CRISPRi phenotypes are moderately correlated (r = 0.68), while CRISPRtune and dCas9 are more strongly correlated (r = 0.92), supporting that MeCP2 is an intermediate repressor capable of tuning gene expression rather than inducing complete repression (**Fig. 5f**). We repeated the growth screen in hTERT-RPE1 cells and results of the screen were consistent across cell types (**Supplementary Fig. 6a-f**). Together, these data highlight the generalizability of CRISPRtune for gene perturbation, allowing for partial repression in contrast to the complete knockdown observed with CRISPRi.

### Orthogonal CRISPR platforms identify genetic dependency partners of MeCP2 and KRAB

Our observations that tethering MeCP2 to gene promoters induces transcriptional tuning provide us with a platform to dissect the genes that are important for MeCP2 repressive activity. Many protein interaction partners of MeCP2 have been well-characterized; however, since MeCP2 binds broadly across the genome, it remains challenging to assess the direct impact of these factors on MeCP2-mediated transcriptional repression. To address this, we performed a knockout arrayed screen of a panel of candidate genes to identify the genetic dependency partners of MeCP2 that are required for its transcriptional tuning activity^66^. We used HEK293T cells with either CRISPRtune or CRISPRi actively silencing CLTA-GFP. We then expressed enhanced Acidaminococcus sp. Cas12a (enAsCas12a) constitutively and corresponding AsCas12a CRISPR RNAs (crRNA) targeting previously hypothesized dependency partners of MeCP2^67^ (**Fig. 6a, b**). AsCas12a CRISPR RNAs were designed using the Humagne library set C and D with two crRNAs per gene on each construct.

**Figure 6.**
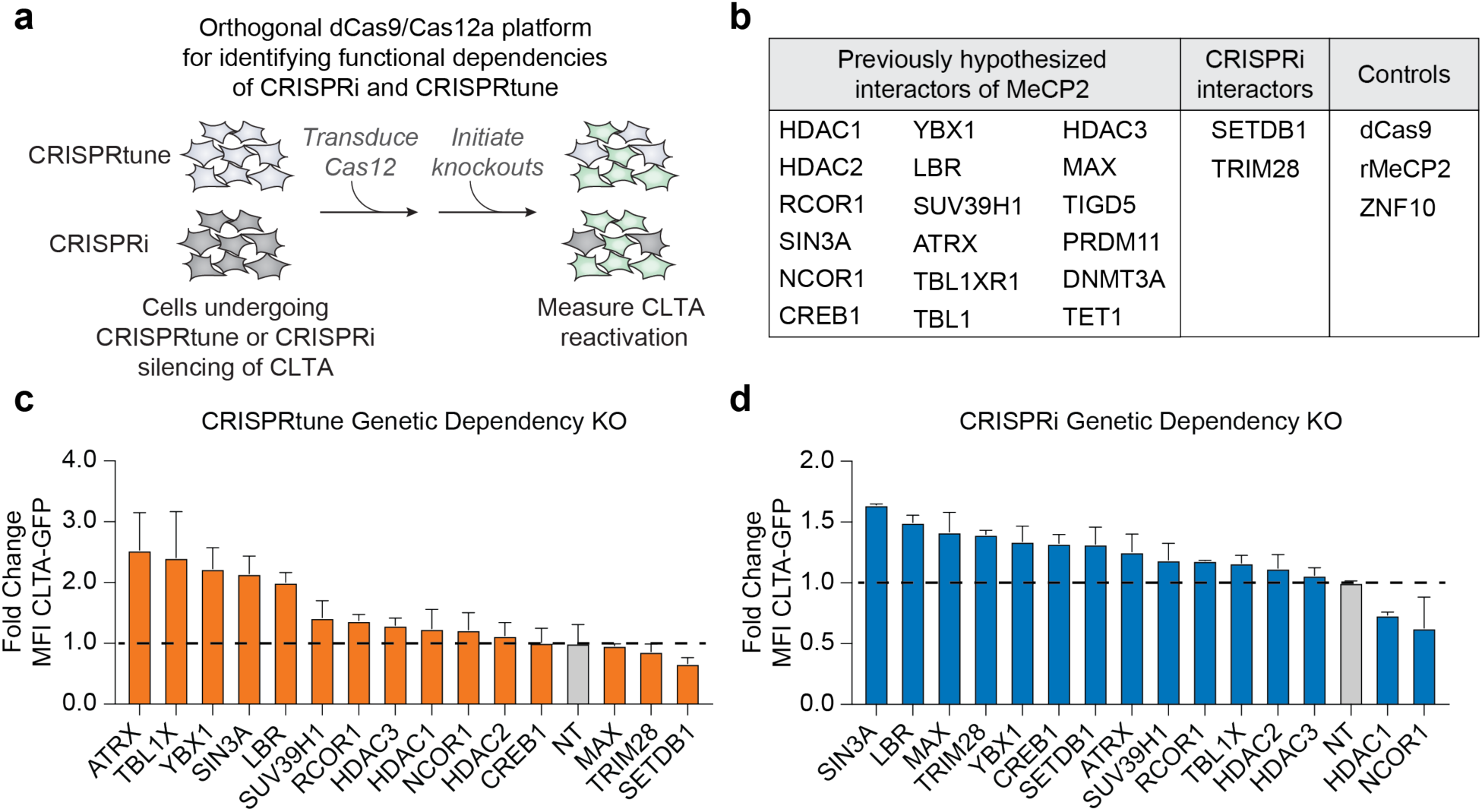
Orthogonal CRISPR platforms for mapping the genetic dependency partners for MeCP2– and CRISPRi-mediated repression. **(a)** A schematic of the arrayed screen workflow. CLTA-GFP HEK293T cells with CRISPRtune or CRISPRi actively silencing CLTA were transduced with enAsCas12a followed by crRNAs for previously hypothesized interactors of MeCP2. **(b)** A table of previously described interactors of MeCP2 tested, as well as validated CRISPRi interactors. dCas9 and rMeCP2 are positive controls for CRISPRtune and ZNF10 is a positive control for CRISPRi. **(c,d)** Quantification of median GFP fluorescence measured 10 days post knockout initiation in cell lines with CRISPRtune (**c**) or CRISPRi (**d**) actively silencing CLTA-GFP in HEK293T cell lines. Data are plotted as fold change relative to a non-targeting Cas12 crRNA. Representative of the top 12 hits from the CRISPRtune cell line and an additional 3 top hits in the CRISPRi cell line. The dashed line is set at the negative non-targeting crRNA control.

As controls, knockout of the dCas9 and MeCP2 components of CRISPRtune leads to loss of repression of CLTA-GFP (**Fig. 6c and Supplementary Fig. 8a**). Notably, knockout of components of the SIN3A complex, including SIN3A, HDAC1, and HDAC2 results in loss of repression of CLTA-GFP. Additionally, knockout of components of the NCoR/SMRT complex, including NCOR1, HDAC3, TBL1X, and RCOR1, also lead to loss of repression. Thus, we conclude that CRISPRtune activity is dependent on the SIN3A and NCoR/SMRT histone deacetylase complexes. Knockout of SETDB1 and TRIM28, known interactors of CRISPRi^51^, results in loss of silencing by CRISPRi and higher CLTA-GFP expression compared to a non-targeting Cas12 crRNA (**Fig. 6d**). Surprisingly, knockout of MAX, a well-known partner of MYC^68,69^, also leads to loss of repression in CRISPRi-expressing lines. Knockouts of SETDB1, TRIM28, and MAX did not lead to loss of repression in CRISPRtune expressing lines, further supporting their role in CRISPRi-mediated repression (**Fig. 6c and Supplementary Fig. 8b**).

Together, these results support a model that KRAB and MeCP2 depend on different pathways that underlie their differing repressive strengths. More broadly, our dCas9-tethering approach, coupled with genome editing, highlights an orthogonal platform for future genome-scale functional genomics experiments to dissect the genetic dependent partners of MeCP2 and other chromatin-regulating proteins.

## DISCUSSION

Here, we present CRISPRtune for transcriptional tuning of endogenous genes in mammalian cells. We show that CRISPRtune is generalizable across thousands of genes, thus highlighting the programmability of MeCP2 for transcriptional repression when targeted to the TSS of promoters. CRISPRtune functions distinctly from KRAB-based CRISPRi as evidenced by their differing epigenetic changes at target gene promoters and their distinct genetic dependency partners for effective transcriptional repression. CRISPRtune functionality is transferable across different cell types, and we apply CRISPRtune for inducing intermediate phenotypes for genome-wide and genome-scale functional genomics screens.

CRISPRtune offers several advantages over existing technologies for tuning gene expression. Compared to RNAi-based repression that is associated with high off-target effects, CRISPRtune relies on the high specificity of CRISPR-dCas9, as supported by our RNA-seq data. Second, we demonstrate that CRISPRtune is scalable across hundreds to thousands of genes, which is a current limitation of degron-based tuning that requires genetic knockins of degron tags at target gene loci. While the dynamic range of repression between dCas9, CRISPRtune, CRISPRtune variants and CRISPRi varies across genes, these editors can be harnessed to elucidate both subtle and significant effects of tuning gene expression. Furthermore, the dynamic range of CRISPRtune can be expanded by introducing RTT-associated variants or using attenuated sgRNAs. We observed that CRISPRtune is more amenable to attenuation using mismatched sgRNAs and results in a more uniformly distributed knockdown of protein expression than CRISPRi. The expanded range of tuning using these strategies with CRISPRtune broadens the applicability of this tool for more precise interrogation of relationships between protein levels and phenotypes.

Our finding that directing MeCP2 to gene promoters induces transcriptional tuning enabled us to dissect the mechanism of MeCP2-dependent transcriptional repression. Mutations in MeCP2 are associated with Rett syndrome, a rare neurodevelopmental disease for which there is no current cure. By engineering the common R306C and P302R Rett syndrome-associated mutations into CRISPRtune, we profiled how defects in MeCP2 transcriptional tuning is due to the failure to remove H3K27ac. In particular, the R306C mutation abolishes CRISPRtune activity, consistent with the mutation failing to recruit the co-repressor complexes NCoR/SMRT^53^. Moreover, genetic knockout of HDAC3 – a core histone deacetylase of NCoR/SMRT – negatively impacts CRISPRtune-dependent repression. In general, our small-scale genetic screens for CRISPRtune genetic dependency factors highlight the utility of dCas9-based tethering to identify genetic dependency factors of disease-associated chromatin factors.

We envision that CRISPRtune will advance targeted gene modulation for use in fundamental research and as a platform for therapeutic applications. For example, several human diseases are due to genetic hypermorphic mutations that can be alleviated therapeutically by reducing gene expression or small molecule-dependent inactivation of protein function. CRISPRtune can be utilized for reducing gene expression to intermediate levels, and its tuning can be further customized when combined with titrating strategies. Furthermore, CRISPRtune is a scalable platform that will be useful in large-scale functional genomic efforts and for targeted single gene perturbations.

## RESOURCE AVAILABILITY

Plasmids encoding MeCP2-dCas9 will be deposited on Addgene upon publication. Bioinformatic datasets for CUT&RUN and RNA-seq experiments are available in the Sequence Read Archive (SRA) of the National Center for Biotechnology Information (NCBI).

Code and analysis pipelines are available at Github: https://github.com/Nunez-Lab

## ACKNOWLEDGMENTS

We thank the Nuñez lab for discussions and feedback on this work. We thank the Chan Zuckerberg San Francisco Biohub Genomics Platform and UCSF CAT (supported by UCSF PBBR, RRP IMIA, and NIH 1S10OD028511-01 grants) for assistance with Illumina sequencing. We thank Jin Chen and Jonathan Weissman for the sgRNA tiling libraries. This project is funded by grants awarded to J.K.N.: the National Institutes of Health (5R35GM155044), The Pew Charitable Trusts, and a University of California Hellman Fellowship. J.K.N. and S.E.C. are Investigators of Biohub, San Francisco. I.J.O., J.I.B., and J.P.L. are funded by the National Science Foundation Graduate Research Fellowship Program. I.J.O. acknowledges funding from the Howard Hughes Medical Institute Gilliam Fellowship and a Mentored Research Award from the University of California, Berkeley. J.I.B. acknowledges funding from the National Institutes of Health Genetic Dissection of Cells and Organisms Training Program (T32GM132022). N.S.D. and L.G.P are funded by postdoctoral fellowships from the California Institute for Regenerative Medicine (EDUC4-12790). S.I.F. acknowledges funding from the Rose Hills Scholarship from the University of California, Berkeley. D.X. is funded by the Weill Neurohub Fellows Program. R.K.P. is funded by graduate fellowships from the University of California Cancer Research Coordinating Committee and the Shurl and Kay Curci Foundation.

## AUTHOR CONTRIBUTIONS

J.I.B., I.J.O, and J.K.N. led the study, designed the experiments and wrote the manuscript with assistance from the co-authors. J.I.B. and I.J.O. performed the majority of the CRISPRtune experiments. P.J.C. performed CUT&RUN experiments, RNA-seq analysis, and ChIP-qPCR experiments. N.S.D. designed and assisted in performing the Cas12 genetic dependency knockout experiments. D.X. prepared RNA-sequencing samples. J.P.L. assisted with computational analysis under the supervision of S.E.C. S.I.F. performed the MeCP2 truncation and HDAC inhibitor experiments. L.G.P. performed the hESC experiments. M.G.H.Z. constructed MeCP2 variants lentiviral plasmids. R.K.P. constructed the human MeCP2-dCas9 plasmid. J.J.M. assisted with designing and performing the enhancer tuning experiments.

## METHODS

### Plasmid DNA generation

The rat MeCP2 sequence was PCR amplified from the Addgene vector #188902 and cloned at the N-terminus of dCas9 within Addgene vector #167981 for CAG-driven transient expression. An XTEN80 linker is positioned between MeCP2 and dCas9. For mRNA expression, the MeCP2-dCas9 fusion was cloned into pALD-CV42 [T7] (Aldevron). For lentiviral expression, the MeCP2-dCas9 fusion was cloned into Addgene vector #188902 for UCOE-SFFV-driven expression or Addgene #188766 for UCOE-EF1a-driven expression. All MeCP2-dCas9 constructs encode a 2x-P2A-BFP linker to measure expression in cells.

The Rett syndrome associated mutants were selected from RettBASE^55^. Mutants from the database were ranked by their frequency, filtered for Rett associated, and then ranked by their clinical severity score from AlphaMissense^56^. The most frequent Rett mutants were selected first, then mutants with a range of clinical severity. The mutations were cloned into the MeCP2-dCas9 construct using Gibson Assembly primers that introduced the point mutations.

### sgRNA design

We utilized the lentiviral pLG1 library vector encodes for expression of an sgRNA from a modified U6 promoter along with expression from an EF1ɑ promoter of a puromycin resistance gene and mCherry separated by a T2A cleavage sequence (Addgene #217306).

Protospacer sequences were ordered as two complementary oligos (IDT) with compatible BstXI and BlpI overhangs and ligated into the lentiviral backbone digested with BstXI and BlpI. Protospacer sequences were chosen based on previous algorithms to predict active CRISPRi sgRNAs targeting gene promoters^70^. A protospacer sequence targeting the *GAL4* gene in *Saccharomyces cerevisiae* was used as non-targeting sgRNA (**Supplementary Table 1**).

### Cell culture and lentivirus production

All cell lines were obtained and authenticated by the UC Berkeley Cell Culture Facility. The CLTA-GFP HEK293T cell lines originated from a previous study^45^. HEK293T were cultured in DMEM (Gibco). K562 cells were cultured in RPMI-1640 (Gibco). h-TERT RPE1 cells were cultured in DMEM/F-12, GlutaMAX. U87-MG cells were cultured in DMEM, supplemented with 1% (v/v) MEM Non-Essential Amino Acids Solution (Gibco) and 1 mM sodium pyruvate (Gibco). All media was supplemented with 10% (v/v) FBS (VWR) and 100 U/mL streptomycin, 100 mg/mL penicillin (Gibco). Cell lines were cultured at 37°C with 5% CO_2_ in tissue culture incubators and routinely confirmed to be negative for mycoplasma.

Human pluripotent stem cells (H9, WA09 Lot #WB68075, WiCell) were cultured in mTeSR Plus (Stem Cell Technologies, 100-0276) and hESC-qualified Matrigel (Corning, 354277)-coated plates following manufacturer’s instructions. Briefly, hESCs-qualified Matrigel was initially pre-diluted in ice-cold DMEM/F12 (Gibco, 10565018) supplemented with 15 mM HEPES (Gibco, 15630080), and plates were incubated at room temperature for an hour before hPSCs were seeded. hPSCs were split every 4-5 days at 70-80% confluency. hPSCs were harvested into cell clumps by incubation for 3 min at 37°C with Gentle Cell Dissociation Reagent (Stem Cell Technologies, 100-0485). Media was refreshed every other day, and cells were cultured at 37°C under normoxia conditions (20% O_2_ and 5% CO_2_).

Lentiviral particles were produced by transfecting standard packaging vectors into HEK293T using TransIT^®^ -LT1 Transfection Reagent (Mirus Bio, MIR2306). Cells were seeded at 4.0×10^5^ cells per well and transfected the next day (1.5 µg lentiviral DNA, 0.1 µg pGag/Pol, 0.1 µg pREV, 0.1 µg pTAT, 0.2 µg pVSVG) with 6 µl TransIT^®^ -LT1 and 250 µl Opti-MEM (Gibco). Media was changed 24 hours post-transfection with complete DMEM. Viral supernatants were harvested 48-60 hours after transfection and filtered using a 0.45 mm PES syringe filter (VWR). Polybrene (8 mg/mL) was used for lentiviral infections.

The dCas9, CRISPRtune, and CRISPRi mRNA were produced using mMESSAGE mMACHINE^TM^ T7 ULTRA Transcription Kit. For each nucleofection reaction in HEK293T cells, 2.0×10^5^ cells were aliquoted, washed with 1 ml of PBS, then resuspended in 20 µl of SF Cell Line Nucleofector^TM^ solution. 2 µg of IVT synthesized editor mRNA was added to the cells resuspended in SF Cell Line Nucleofector^TM^ solution and transferred to a Lonza Nucleocuvette^TM^. The Nucleofector^TM^ program CM-130 was used on the 4D-Nucleofector ^TM^ System. Following nucleofection, cells were resuspended in 80 µl of DMEM and transferred into a well of a 24-well plate containing 400 µl of pre-warmed culture medium. For plasmid nucleofection in H9 hESC cells, cells were collected in single cells using StemPro Accutase (Gibco, A1110501) and incubated for 3 minutes at 37°C. 5.0×10^5^ cells were resuspend in 20 µl of P3 Cell Line Nucleofector^TM^ solution and a mix of sgRNA and editor plasmids (5 µg of DNA in total with a 3:1 ratio sgRNA:editor). The Nucleofector^TM^ program CB-150 was used on the 4D-Nucleofector^TM^ System. After the pulse, cells were seeded in mTeSR Plus supplemented with Cloner2 (Stem Cell Technologies, 100-0691). After 24 hours, media was changed to regular mTeSR Plus until data collection at 72 hours post-nucleofection.

### Antibody staining and flow cytometry

Protein expression was assessed by cell surface antibody staining of live cells. Cells were incubated with diluted antibody (**Supplementary Table 2**) for 30-60 minutes in the dark at room temperature, washed once with PBS and measured on the BD FACSymphony^TM^ A1 Cell Analyzer (BD Biosciences). For CLTA-GFP HEK293T reporter lines, CLTA-GFP protein expression was assessed directly by flow cytometry. Data analysis was performed using FlowJo^TM^ (v10.10).

### Genetic dependency experiments

To generate the Cas12 cell lines, tuned or silenced CLTA-GFP HEK293T and hTERT-RPE1 were sorted for CRISPRtune or CRISPRi expressing cells, respectively. To introduce the Cas12, mCherry or bleomycin resistance was cloned into the C-terminus of EnAsCas12a lentiviral vector pRDA_174 (Addgene #136476) separated by T2A. Cells were sorted for mCherry or selected for Cas12 using 500 µg/mL zeocin (ThermoFisher, R25001) for 10 days.

To generate the Cas12 knockout crRNAs, crRNA sequences were cloned from Humagne library C and D^67^. Genetic dependency factors were selected from the STRING database^66^. crRNA sequences were ordered as complementary sequences from IDT, annealed, and then ligated into PRDA_052 EnAsCas12a sgRNA vector (Addgene # 136474) using AgeI and EcoRI restriction sites. Lentivirus was prepared by transfection with standard lentiviral packaging vectors along with Cas12 knockout crRNA plasmid using TransIT^®^ -LT1 Transfection Reagent (Mirus Bio, MIR2306) and harvested 2 days post transfection and filtered using a 0.45 µm filter.

Cas12 crRNA lentivirus was added to the Cas12 expressing cells and selected for using 2 µg/mL puromycin. Cells were split every 2 days and GFP expression was measured on day 7 and day 10 by flow cytometry on the BD FACSymphony^TM^ A1 Cell Analyzer (BD Biosciences). Protein expression of CD55 in hTERT-RPE1 was determined by antibody staining using the method described above. Data analysis was performed using FlowJo^TM^ (v10.10).

### Attenuating sgRNA experiments

Attenuating sgRNAs targeting *CD55* were selected from a published dataset predicted to have differential CRISPRi activity^24^. hTERT-RPE1 cells were seeded in 24-well plates and transduced with lentivirus carrying each sgRNA into cell lines stably expressing dCas9, CRISPRi, or CRISPRtune in biological triplicates. After transduction, cells were selected with puromycin (1 µg/mL) until >90% of cells were stably expressing sgRNAs. Following selection, cells were stained with anti-CD55 antibody and analyzed by flow cytometry. Median fluorescence intensity of CD55 was quantified for each sgRNA to assess repression strength.

### Enhancer targeting experiment and qPCR quantification

K562 cells stably expressing CRISPRi or CRISPRtune were spinfected with lentivirus (1000 × g for 2 hours at 33 °C) encoding sgRNAs (mCherry) targeting the non-targeting control (NT), TSS, or enhancer region of *GATA1* using previously validated sgRNA sequences. 2 days post-transduction, cells were sorted for mCherry positive cells. Cells were grown for 9 days post-transduction and total RNA was extracted using RNeasy Mini Kits (Qiagen) following the manufacturer’s protocol. cDNA was synthesized from 250 ng RNA per sample using the ProtoScript II First Strand cDNA Synthesis Kit (NEB). qPCR was performed using iTaq Universal SYBR Green Supermix (Bio-Rad) and relative mRNA was calculated using ΔΔCq, normalizing each sample to its parenteral cell line (**Supplementary Table 3**).

### Genome-scale sgRNA tiling screen

The sgRNA tiling library was designed as described in detail previously. Briefly, we selected NGG-spanning targets across a ± 2 kb region relative to the TSS of 1,869 genes. The final library consisted of 472,632 unique sgRNAs of which 37,900 were non-targeting negative control sgRNAs (8% of total). The pooled library was packaged in lentiviral particles and infected into 720×10^6^ K562 cells by spinfection for 2 hours at 33°C with 8 µg/mL polybrene. Then, the population was split into two technical replicates and cultured in spinner flasks at 37°C with 5% CO_2_. Two days post-transduction, 30% of sgRNA-expressing cells was detected (mCherry marker fluorescence) by flow cytometry. Cells were treated with puromycin until the mCherry-positive cells reached ∼94-95% of the total population, which served as the initial time point (T*initial*). Cells were passaged daily with cell counting, measurement of viability, and mCherry-positive percentage. Cells were collected at T*final* when the cells reached ∼11 population doublings. Thereafter, genomic DNA extraction, sgRNA identification by Illumina sequencing, and downstream processing and phenotype score calculations were performed as described previously^71^.

### Essential gene screen

For the growth-based screen targeting essential genes in K562 cells, an sgRNA library was designed based on a previously published genome-wide CRISPRi screen in K562s and the DepMap CRISPR-inferred common essential genes^60,64^. The library consists of 8,086 sgRNAs targeting 1,838 essential genes, along with non-essential and non-targeting control sgRNAs. Each essential gene is targeted by four sgRNAs, selected from previously published sgRNA libraries^39,71^. A list of protospacer sequences included in the essential-gene library is included in **Supplementary Table 1.**

sgRNA sequences were cloned into an expression plasmid obtained from Addgene (ID#217306), which includes a U6 Pol III promoter and a T2A-mCherry reporter to measure transduction efficiency. The sgRNA oligos were synthesized as a pooled library (Twist Bioscience) with the structure: 5’ – PCR adaptor – CCACCTTGTTGG – protospacer sequence – GCTAAGC – PCR adaptor – 3’. Oligo pools were PCR-amplified, digested with BstXI/BlpI, gel extracted, and ligated into the sgRNA lentiviral vector. The resulting plasmid library was transformed for amplification and validated by next-generation sequencing to confirm library representation and uniformity.

To conduct the screen, K562 cells stably expressing a dCas9 fusion protein were transduced with the lentiviral sgRNA library via spinfection (1000 × g for 2 hours at 33 °C) with polybrene (8 µg/mL, Sigma-Aldrich). Infection efficiency was assessed two days post-infection by flow cytometry, aiming for 20%–30% mCherry-positive (sgRNA-expressing) cells. Screens were performed with two technical replicates, maintaining a minimum representation of 1,000 cells per sgRNA throughout the experiment. Two days post-transduction, puromycin was added to the media until ∼90% of cells were sgRNA-positive, as indicated by mCherry fluorescence. On day 6 post-transduction, cell pellets were collected and frozen for each replicate to represent the initial time point (T0). Cell doublings were tracked throughout the experiment, and final samples (Tfinal) were collected after approximately 12 cell doublings.

DNA libraries for T0 and Tfinal were prepared for deep sequencing following previously described protocols^24^. Genomic DNA was extracted from cell pellets using the NucleoSpin Blood L kit (Macherey–Nagel). The gDNA was directly PCR-amplified (23 cycles) using NEBNext Ultra II Q5 PCR Master Mix (NEB) to append Illumina adapters and unique sample indices. Libraries were sequenced on a NovaSeqX platform with a 19 bp Read 1 and a 5 bp Index 1.

### RNA-sequencing

hTERT RPE-1 cells were maintained for 10 days post-transduction of dCas9, CRISPRtune, and CRISPRi targeting *CD55*. Cells were centrifuged at 500 x g for 5 minutes and washed with PBS. Total RNA was extracted using the RNeasy Mini Kit (QIAGEN). Poly(A) mRNA was enriched using poly(A) mRNA Magnetic Isolation Module (New England Biolabs). Library preparations were carried out using NEBNext^®^ Ultra^TM^ II Directional RNA Library Prep Kit for Illumina (New England Biolabs), starting with 1000 ng total RNA. Final libraries were assessed using an Agilent 2100 TapeStation system (Agilent), and sequenced as paired end 50 base pair reads on a NovaSeq X (Illumina).

Reads were trimmed using cutadapt (version 4.9) filtering empty resulting reads (-m 1) and specifying both forward (-a AGATCGGAAGAGCACACGTCTGAACTCCAGTCA) and reverse (-A AGATCGGAAGAGCGTCGTGTAGGGAAAGAGTGT) adapters^72^. The resulting reads were aligned using Kallisto (version 0.51.1) to the human cDNA transcriptome (GRCh38 release 114)^73^. RNA isoforms were aggregated to genes using tximport (version 1.36.0)^74^. Protein-coding genes identified and filtered via biomaRt (version 2.64.0) using R (version 4.5.0)^75^. Differential gene expression analysis was performed using DESeq2 (version 1.48.1)^73^. All downstream analyses were performed on python (version 3.11.7) using polars (version 1.27.1).

## CUT&RUN

CUT&RUN libraries were generated using the CUTANA™ ChIC/CUT&RUN kit following the manufacturer’s manual. 550,000 cells were collected per experimental condition. The cells were permeabilized in 0.05% digitonin (Epicypher 21-1004). Antibody incubation was performed overnight (18-20 hours) at 4°C. For H3K4me3 (Epicypher 13-0060), H3K27ac (Epicypher 13-0059), and IgG (Epicypher 13-0042) antibody incubations, 1 µL (0.5 µg) was added per reaction. For H3K9me3 (Abcam ab176916), 1.5 µL (1.93 µg) was added per reaction. Libraries were generated using NEBNext^®^ Ultra^TM^ II DNA Library Prep Kit for Illumina (NEB E7645L). For K562 cells, QB3 Truseq plate WhiteV4 primers were used for barcoding while for h-TERT RPE1 cells, NEBNext^®^ Multiplex Oligos for Illumina® (NEB E6440S) were used. For H3K4me3, H3K9me3 and H3K27ac antibody conditions, 3 ng of input was PCR-amplified for 8 cycles. For IgG antibody conditions, 1 ng of input was PCR-amplified for 10 cycles. Samples were PCR-cleaned up and eluted in 1X TE according to the protocol. Samples were quantified using an Agilent 4150 TapeStation device using D1000 DNA ScreenTape (Agilent 5067-5582), and were equally pooled to a final concentration of 19.22 nM for K562 cells and 5.28 nM for hTERT-RPE1 cells. The prepared libraries were sequenced pair-end 50 bp reads on an Illumina NovaSeq X 1.5B 1 lane (0.8 billion clusters) for K562 cells and NovaSeq X 10B 1 Lanes (1.3 billion clusters) for hTERT-RPE1 cells.

Reads were trimmed using cutadapt (version 4.9) filtering empty resulting reads (-m 1) and specifying both forward (-a AGATCGGAAGAGCACACGTCTGAACTCCAGTCA) and reverse (-A AGATCGGAAGAGCGTCGTGTAGGGAAAGAGTGT) adapters^72^. The resulting reads were aligned using Bowtie2 (version 2.5.2) to the human genome index (GRCh38; Langmead, “Bowtie 2 indexes”)^74^. Mate pair information was filled, reads were deduplicated, and DAC blacklist sites were removed^76,77^ using Samtools (version 1.19.2)^75^. Bedgraph files were generated using deepTools bamCoverage (version 3.5.4)^78^. IGV was used to generate coverage tracts (version 2.19.4)^79^.

Differential peak analysis was performed by calling broadpeaks with macs3 callpeak (--broad --broad-cutoff 0.1) (version 3.0.3)^80^. Next, DiffBind (version 3.12.0) was used to perform differential peak analysis in R (version 4.3.3) using dba.count (minOverlap=1, summits=250), dba.normalize, dba.analyze (method=DBA_ALL_METHODS), and dba.report (method = DBA_EDGER, contrast = 1, th = 1, bUsePval = FALSE)^81,82^. Volcano plots were made in python (version 3.11.7) using matplotlib (version 3.10.3)^83^.

### Chromatin Immunoprecipitation quantitative polymerase chain reaction

20-30×10^6^ cells were harvested and crosslinked in 1% formaldehyde in DPBS for 10 min at room temperature then quenched to 200 mM glycine for 10 min at room temperature. Pellets were lysed in 1 mL lysis buffer (5mM PIPES pH 8.0, 85mM KCl, 1% Igepal, and 1X Protease Inhibitor) per 10^6^ cells on ice for 10 min. Pellets were flash frozen in liquid nitrogen and stored in -80 °C. Next, nuclei were isolated by centrifuging at 1000 x g at 4 °C for 5 min. The supernatant was aspirated and the nuclei were lysed in 300 µL of nuclei lysis buffer (50 mM Tris pH 8.0, 10 mM EDTA, 1% SDS, 1X protease inhibitor) for 10 minutes on ice. Chromatin shearing was performed using a Diagenode Bioruptor® Pico sonication device in 1.5 mL Bioruptor® Pico Microtubes (10 cycles 30 sec on 30 sec off). The sonication samples were centrifuged at 13,000 RPM at 4 °C, and chromatin supernatant was collected and diluted 4-fold in IP dilution buffer (50 mM Tris pH 7.5, 150 mM NaCl, 1 mM EDTA, 1% Igepal, 0.25% Na-DOC, 1X protease inhibitor). 25 µL of chromatin was reverse crosslinked and analyzed on a 2% agarose gel to verify sonication. Glycerol was added to 5% to the chromatin, quantified using the Qubit™ dsDNA Quantification Assay Kits on a Denovix® DS-11 FX. Chromatin was flash frozen in liquid nitrogen and stored in -80 °C. 30 µg of chromatin was diluted in IP dilution buffer to a final volume of 1 mL. A 4% input fraction was collected and frozen for downstream quantification. Immunoprecipitation was performed overnight at 4 °C using 2 µL of antibody (Cell Signaling Technology 9649S, 8173S). 50 µL of Protein A/G magnetic beads (Thermo Fisher 88803) per sample were washed as follows: 1) place beads in a magnetic tube rack and aspirate supernatant, 2) wash at least twice in 500 µL IP dilution buffer, 3) resuspend in 50 µL of IP dilution buffer, 4) add 50 µL of washed beads into ChIP sample. The samples were rotated for 4 hours at 4 °C. The samples were washed as follows: 1) once in 1 mL of IP wash buffer 1 (20 mM Tris pH 8.0, 2 mM EDTA, 50 mM KCl, 1% Triton X-100, 0.1% SDS), 2) twice in 1 mL High salt buffer (20 mM Tris pH 8.0, 2 mM EDTA, 500 mM NaCl, 1% Triton X-100, 0.01% SDS), 3) once in 1 mL IP wash buffer 2 (10 mM Tris pH 8.0, 1 mM EDTA, 0.25 M LiCl, 1% Igepal, 1% Na-DOC), 4) twice in TE Buffer (10 mM Tris pH 8.0, 1 mM EDTA). The samples were eluted in 100 µL Elution buffer (50 mM NaHCO3, 1% SDS) under constant agitation for 1 hour at 65 °C on a thermomixer then reverse crosslinked. For the immunoprecipitation products, 5 µL of 5 M NaCl was added to each sample. The inputs were diluted to 100 µL of IP dilution buffer, then 3 µL of 5 M NaCl was added. 1 µL of RNAse A was added to each sample and input and incubated for 1 hour at 37 °C. Next, 2.25 µL of proteinase K was added and incubated overnight at 63 °C. Samples were purified using the Qiagen QIAquick PCR Purification Kit (Qiagen 28104) and eluted in two rounds of 20 µL of buffer EB.

The ChIP products and inputs were quantified using the Qubit™ dsDNA Quantification Assay Kits on a Denovix® DS-11 FX to ensure there was sufficient pulldown. DNA was amplified using a Biorad CFX Opus 96 Real-Time PCR system using iTaq™ Universal SYBR® Green Supermix (Biorad 1725121) using 2 µL of sample following manufacturer’s recommendation. Fold enrichment normalization was performed to a negative control site (ZNF510).

### Trichostatin A Inhibitor Experiments

To conduct the TSA inhibitor experiments, hTERT-RPE1 cells were plated at 2×10⁵ cells per well in 24-well plates containing a total volume of 500 μL solution per well. Each experimental condition (WT, CRi, MeCP2t, and CRISPRtune) was performed in triplicate. Cells were treated with TSA at final concentrations of 6.25, 12.5, 25, 50, or 100 nM. Also included was an untreated (no DMSO) and vehicle (0 nM TSA in DMSO) control conditions.

TSA was purchased from Selleck Chemicals (S1045) and resuspended in DMSO (Sigma Aldrich RNBN8586). A 10 mM TSA stock solution was diluted 1:100 in DMSO to generate a 100 μM working stock. Two-fold serial dilutions were prepared to obtain 100, 50, 25, 12.5, and 6.25 μM working solutions. A vehicle control containing DMSO alone was also prepared with 10μL of DMSO and the rest media. For each condition, 10 μL of the appropriate working solution was added to 5 mL of cell media respectively. Aliquots of 250 μL of the prepared solution were then added to each well containing 250 μL of previously added cells with media to achieve final TSA concentrations of 6.25, 12.5, 25, 50, or 100 nM and a total volume of 500μL per well. The final DMSO concentration in all treated and vehicle control wells was maintained below 0.05%.

Cells were incubated with TSA for 48 hours before harvesting for flow cytometry. Following treatment, cells were stained with an APC-human CD55 antibody and analyzed by flow cytometry. Data was analyzed using FlowJo^TM^ (v10.10).

## SUPPLEMENTARY FIGURES

**Supplementary Figure 1.**
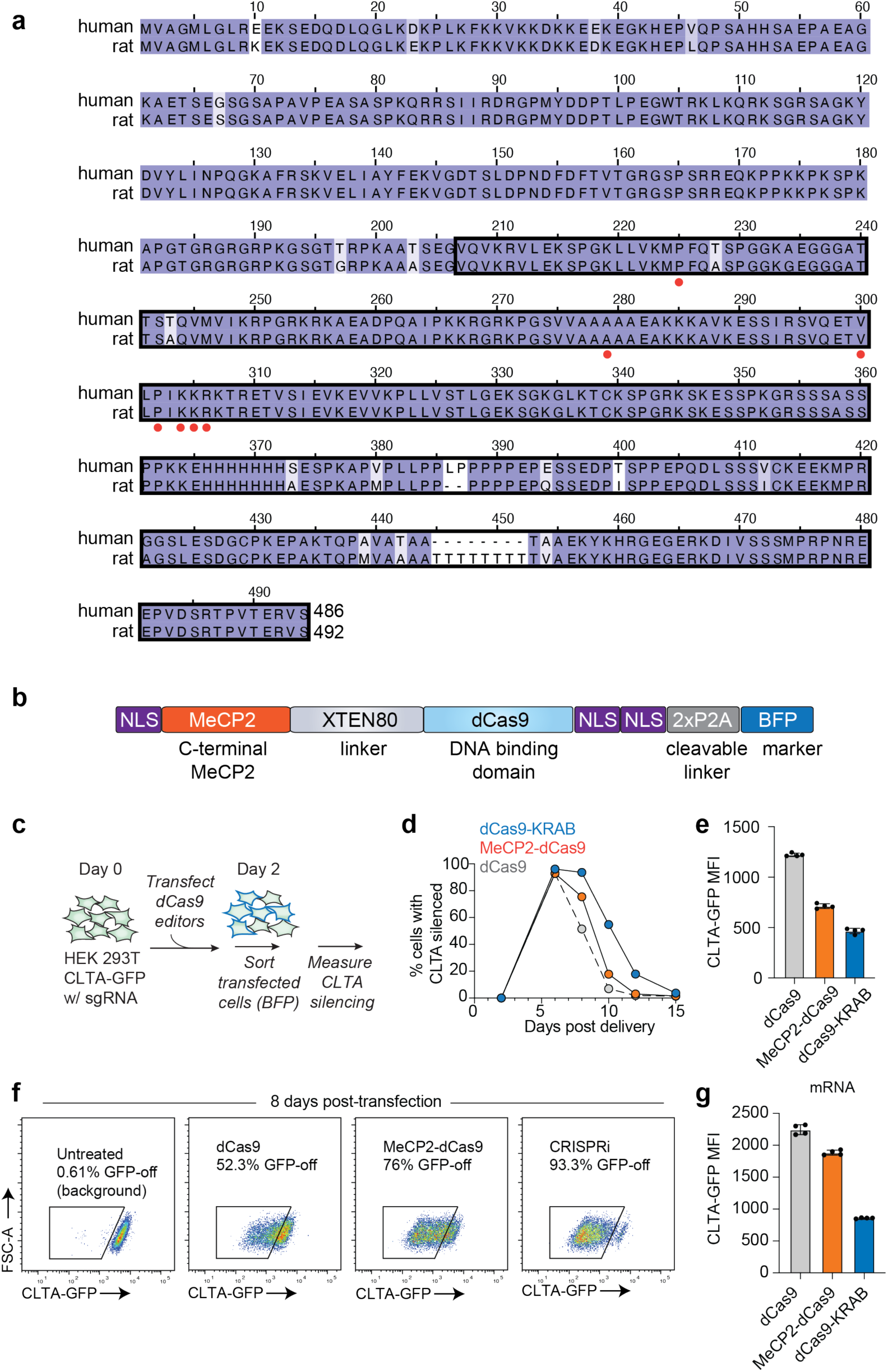
Programmable tuning of endogenous genes with transient expression of CRISPRtune. **(a)** An alignment of the human and rat orthologs of MeCP2. The region in the black box was used in the MeCP2-dCas9 fusion (CRISPRtune). Red dots indicate locations of Rett Syndrome mutations tested in Figures 3A-3D. **(b)** A schematic of the CRISPRtune construct design used in the study. **(c)** A schematic of the workflow testing transfections of dCas9-KRAB, MeCP2-dCas9, and dCas9 in CLTA-GFP HEK293T cells constitutively expressing an sgRNA targeting *CLTA*. **(d)** A time course of CLTA silencing following transfection of dCas9-KRAB (blue), MeCP2-dCas9 (orange), and dCas9 (gray). **(e)** Quantification of CLTA-GFP median fluorescence intensity 8 days post-transfection. **(f)** Representative flow cytometry plots of CLTA-GFP levels 8 days post-transfection. **(g)** Quantification of CLTA-GFP median fluorescence intensity 2 days post-mRNA nucleofection.

**Supplementary Figure 2.**
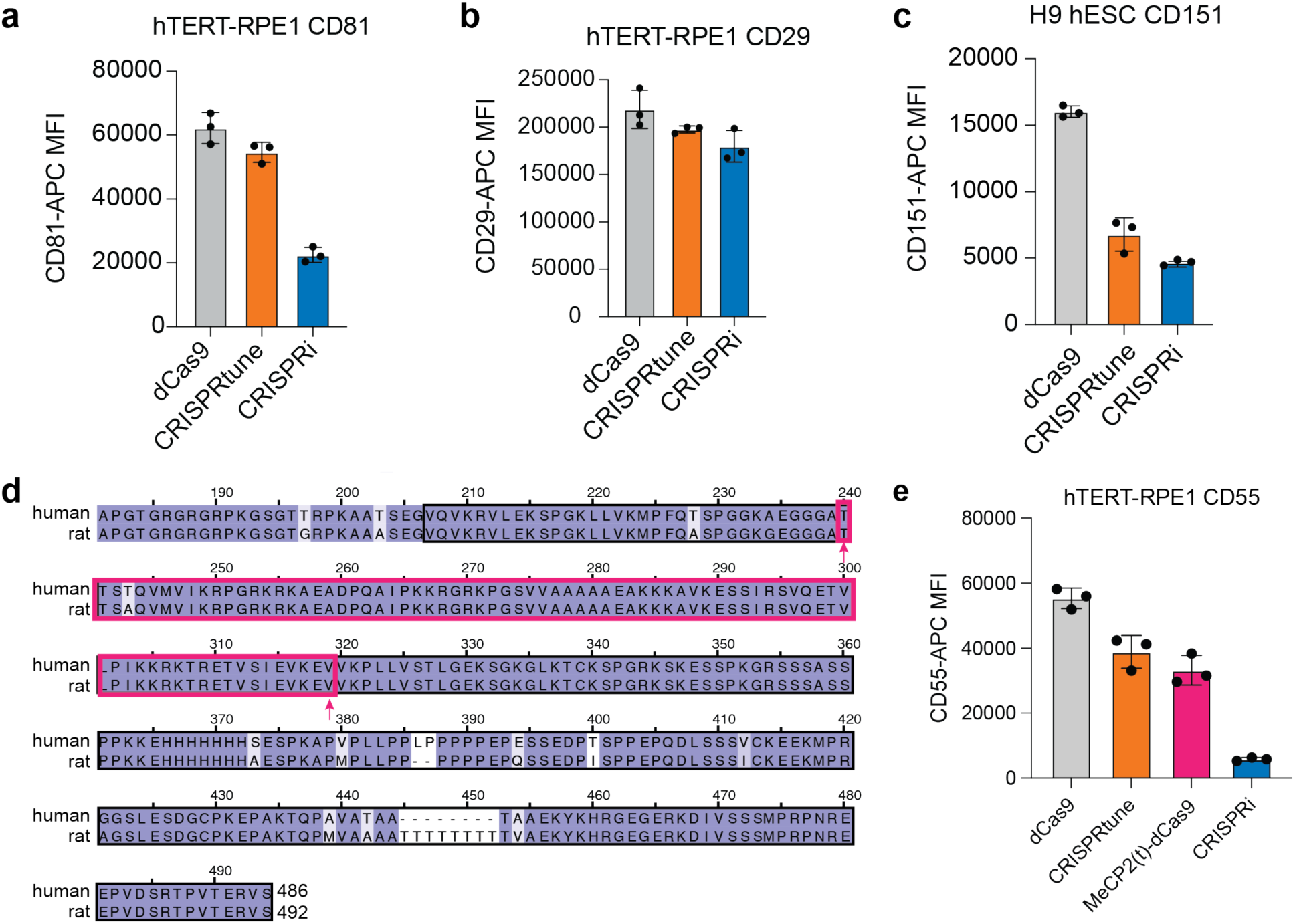
CRISPRtune is a transcriptional tuner across genes and cell types. (**a, b**) Quantification of CD81 **(a)** and CD29 levels **(b)** in hTERT-RPE1 levels following lentiviral transduction of dCas9, CRISPRtune, and CRISPRi and an sgRNA targeting *CD81* and *CD29*. For CD81, CRISPRtune repression is intermediate between dCas9 (*p* = .0851) and CRISPRi (*p* = .0001). For CD29, CRISPRtune repression is intermediate between dCas9 (*p* = .1475) and (*p* = .1425). **(c)** Quantification of CD151 levels in H9 hESC following plasmid nucleofection of each editor with a CD151 sgRNA. CRISPRtune repression is intermediate between dCas9 (*p* = .0003) and CRISPRi (*p* = .0456). **(d)** An amino acid alignment of rat and human MeCP2. The region included in the truncation is outlined in pink, while the original MeCP2 fusion used in CRISPRtune is outlined in black. **(e)** Quantification of CD55 levels in hTERT-RPE1 cells constitutively expressing a CD55 sgRNA following lentiviral transduction of dCas9, CRISPRtune, dCas9-MeCP2(t) (MeCP2 truncation from (**d**)), and CRISPRi. MeCP2(t) – dCas9 represses *CD55* to similar levels as CRISPRtune (*p* = .2199) but is significantly different from dCas9 (*p* = .0023) and CRISPRi (*p* = .0005).

**Supplementary Figure 3.**
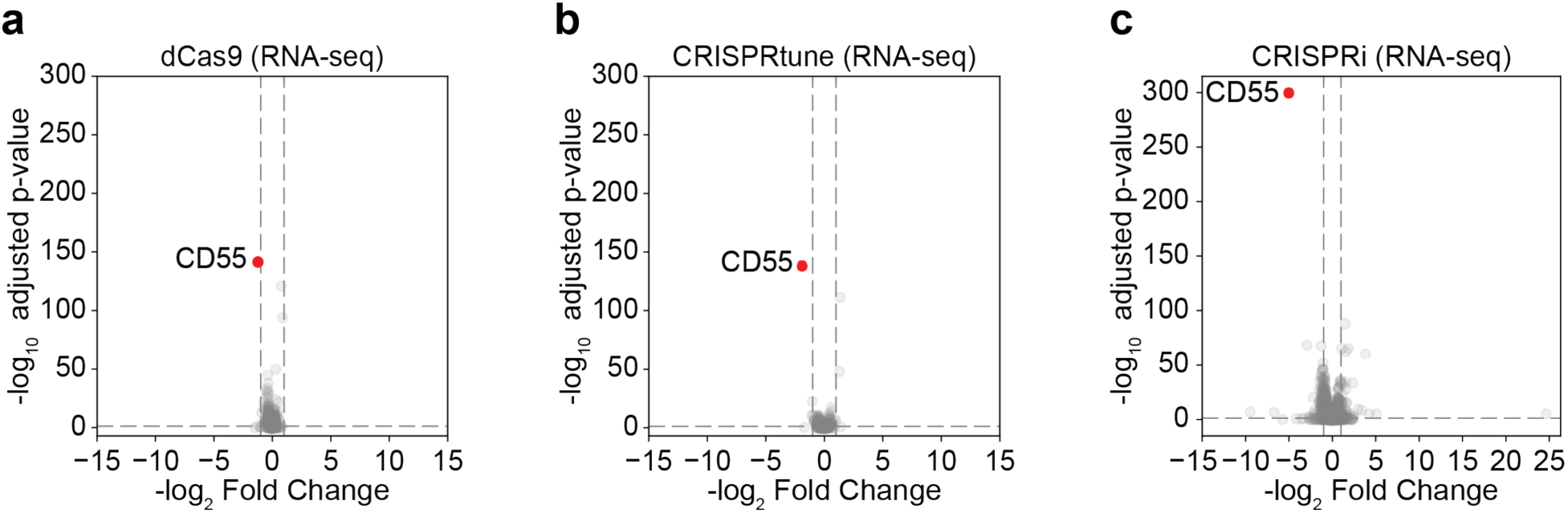
Transcriptional tuning by CRISPRtune. **(a-c)** Volcano plots comparing RNA-seq of dCas9 (**A**), CRISPRtune (**B**), and CRISPRi (**C**) targeting *CD55*. Red dots represent *CD55* transcripts.

**Supplementary Figure 4.**
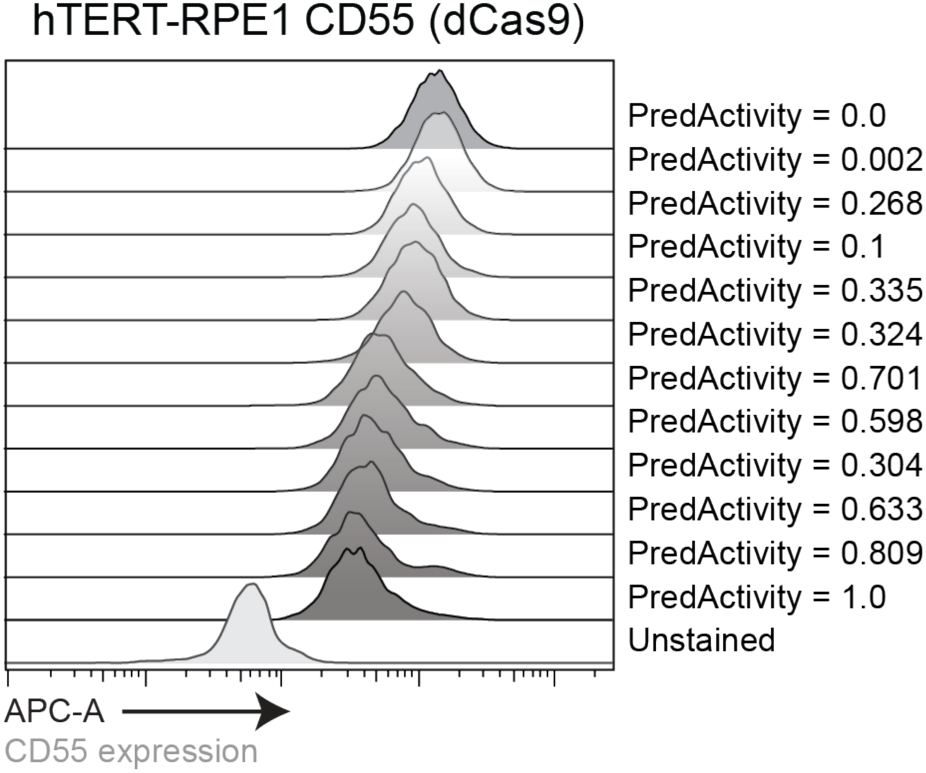
Attenuated sgRNAs titrate dCas9 gene repression. A histogram plot of flow cytometry data from hTERT-RPE1 cells that stably express dCas9 and the corresponding attenuated sgRNAs.

**Supplementary Figure 5.**
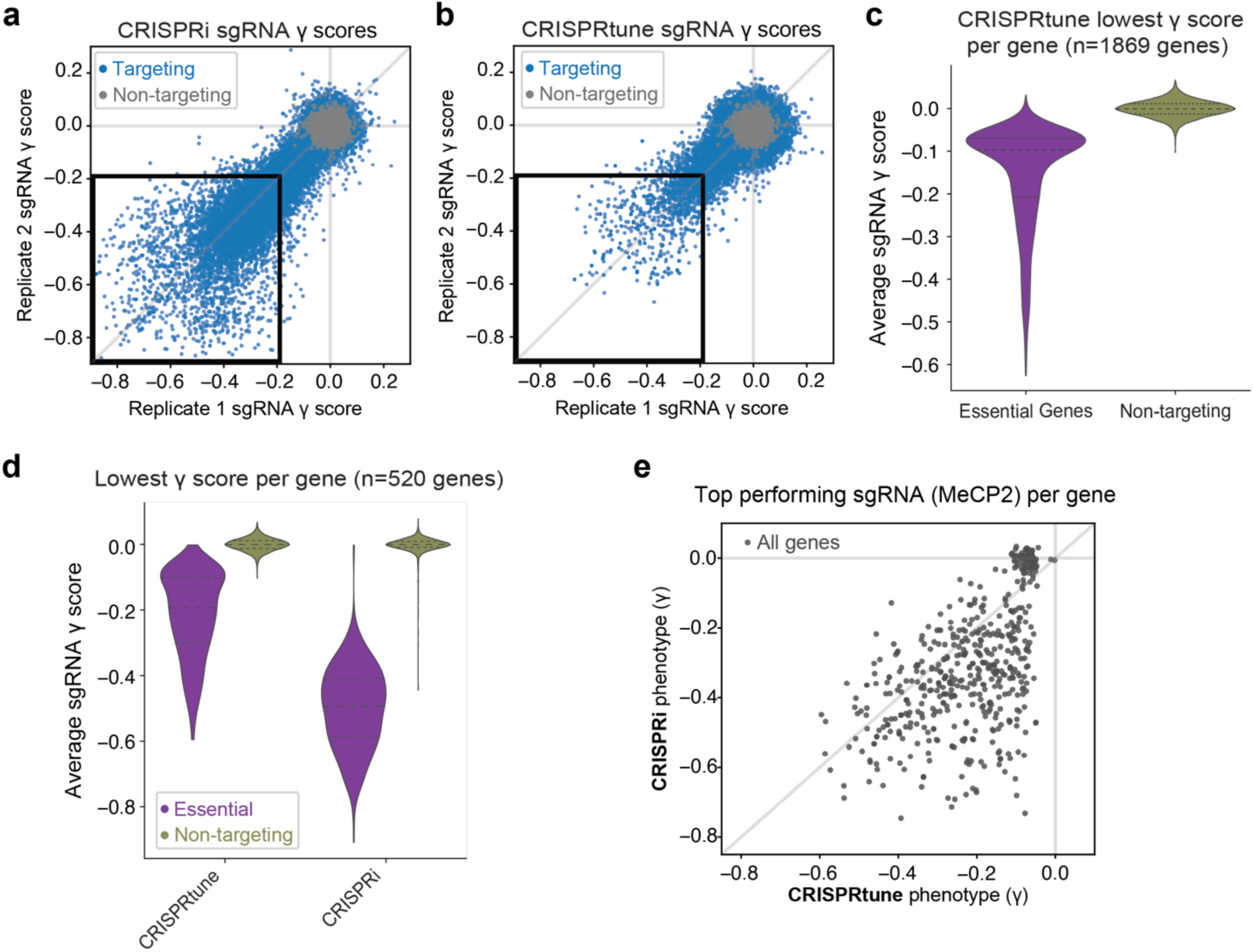
CRISPRtune modulates gene expression in a pooled sgRNA tiling screen. **(a,b)** Scatterplots showing replicate correlations of sgRNA phenotype scores (γ) from pooled tiling gene screens performed in K562 cells expressing CRISPRi (**a**) and CRISPRtune (**b**). Each dot represents an individual sgRNA targeting essential genes (blue) or non-targeting controls (gray). **(c)** Violin plots of lowest gene phenotype scores for CRISPRtune for 1,869 essential genes and non-targeting controls. **(d)** Violin plots of lowest gene phenotype scores for CRISPRtune and CRISPRi for 520 essential genes and non-targeting controls. **(e)** Scatter plot showing phenotype scores for the best performing sgRNA for CRISPRtune and CRISPRi respectively.

**Supplementary Figure 6.**
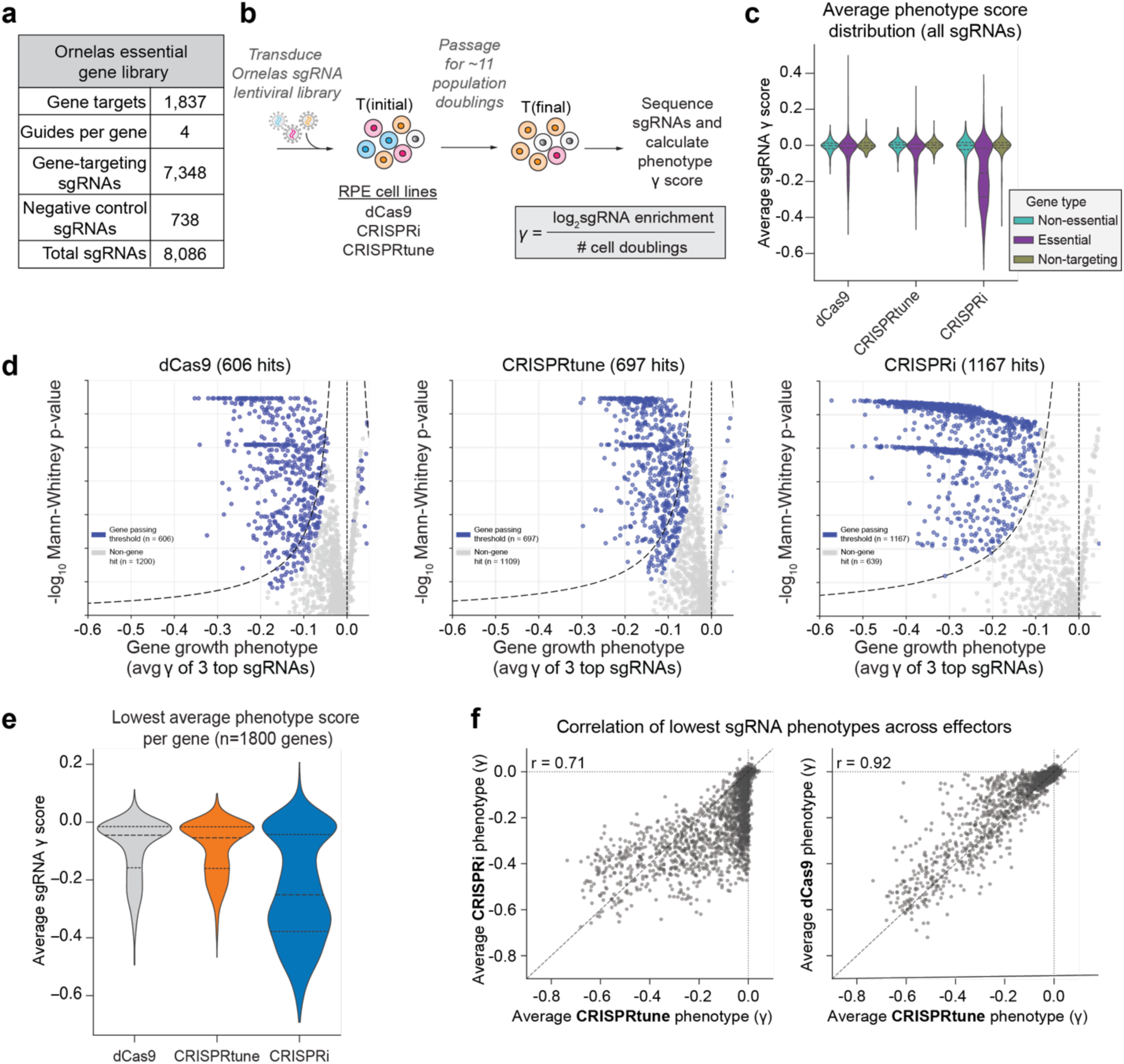
CRISPRtune modulates essential gene phenotypes across cell types. **(a)** A summary of the number of genes and sgRNAs per indicated category in the Ornelas essential gene library. **(b)** A schematic of the essential gene library screen workflow in hTERT-RPE1 (RPE) cells. **(c)** A violin plot of the average sgRNA phenotype score (γ) for dCas9, CRISPRtune, and CRISPRi. Populations are grouped by sgRNAs for nonessential genes (teal), essential genes (purple), and non-targeting (yellow). **(d)** Volcano plots comparing the average gene growth phenotype of 3 top sgRNAs (x axis) and the -log10 Mann-Whitney p-value. Blue dots represent genes passing the threshold of significance and gray dots represent non-gene hits. **(e)** A violin plot of the lowest average phenotype scores for each gene (left). A Venn diagram comparing the number of gene hits (γ < 0.1) for dCas9 (gray) and CRISPRtune (orange). **(f)** A comparison of the lowest sgRNA phenotype for each sgRNA between CRISPRtune and both CRISPRi (left) and dCas9 (right).

**Supplementary Figure 7.**
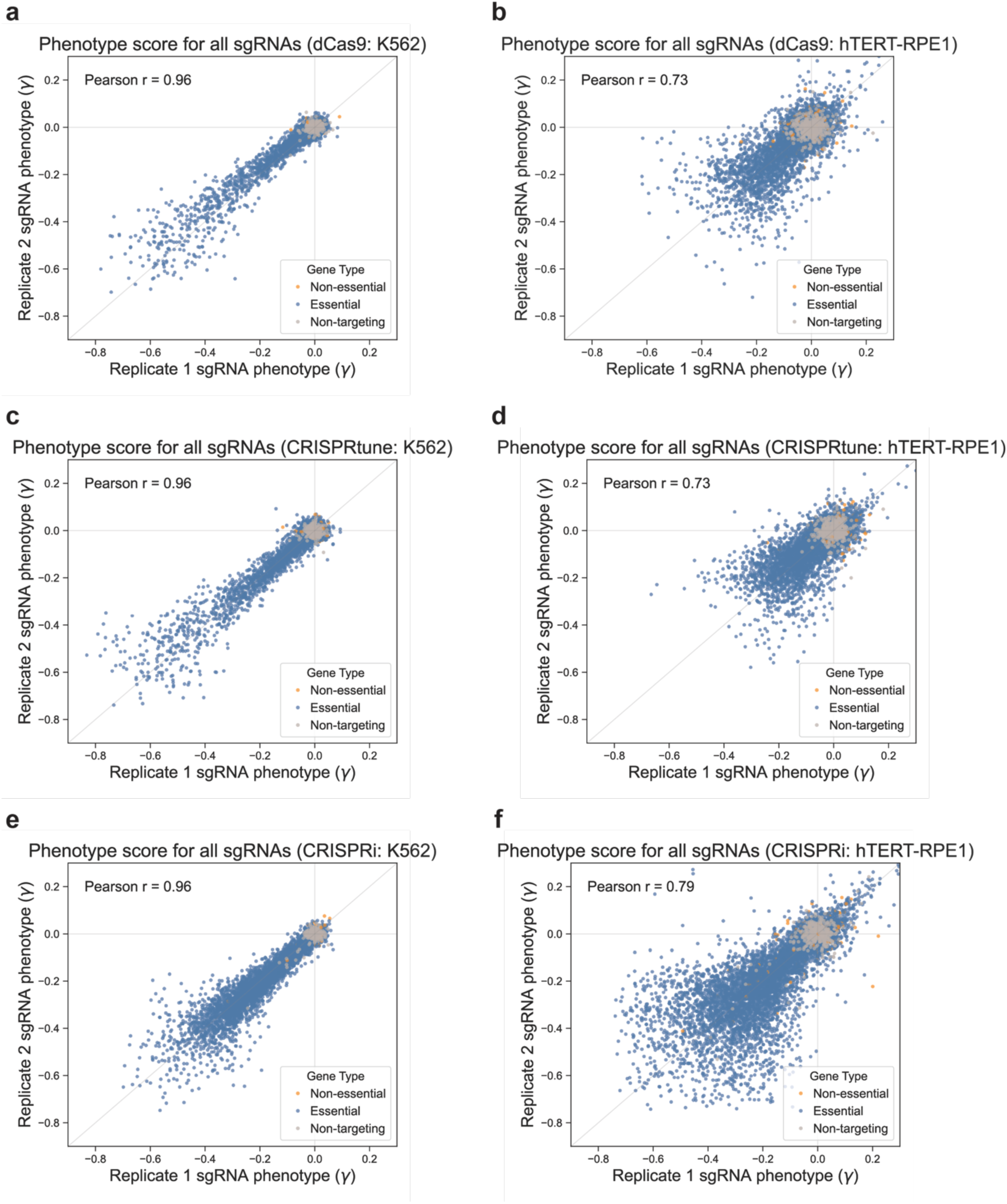
Essential gene screen phenotypes are consistent across replicates. **(a-f)** A comparison of phenotype scores for all sgRNAs in the essential gene screen across replicate 1 (x axis) and replicate 2 (y axis). **(a)** dCas9 (K562 screen, Figure 5) **(b)** dCas9 (hTERT-RPE1 screen, Supplementary Figure 5) **(c)** CRISPRtune (K562 screen) **(d)** CRISPRtune (hTERT-RPE1 screen) **(e)** CRISPRi (K562 screen) **(f)** CRISPRi (hTERT-RPE1 screen)

**Supplementary Figure 8.**
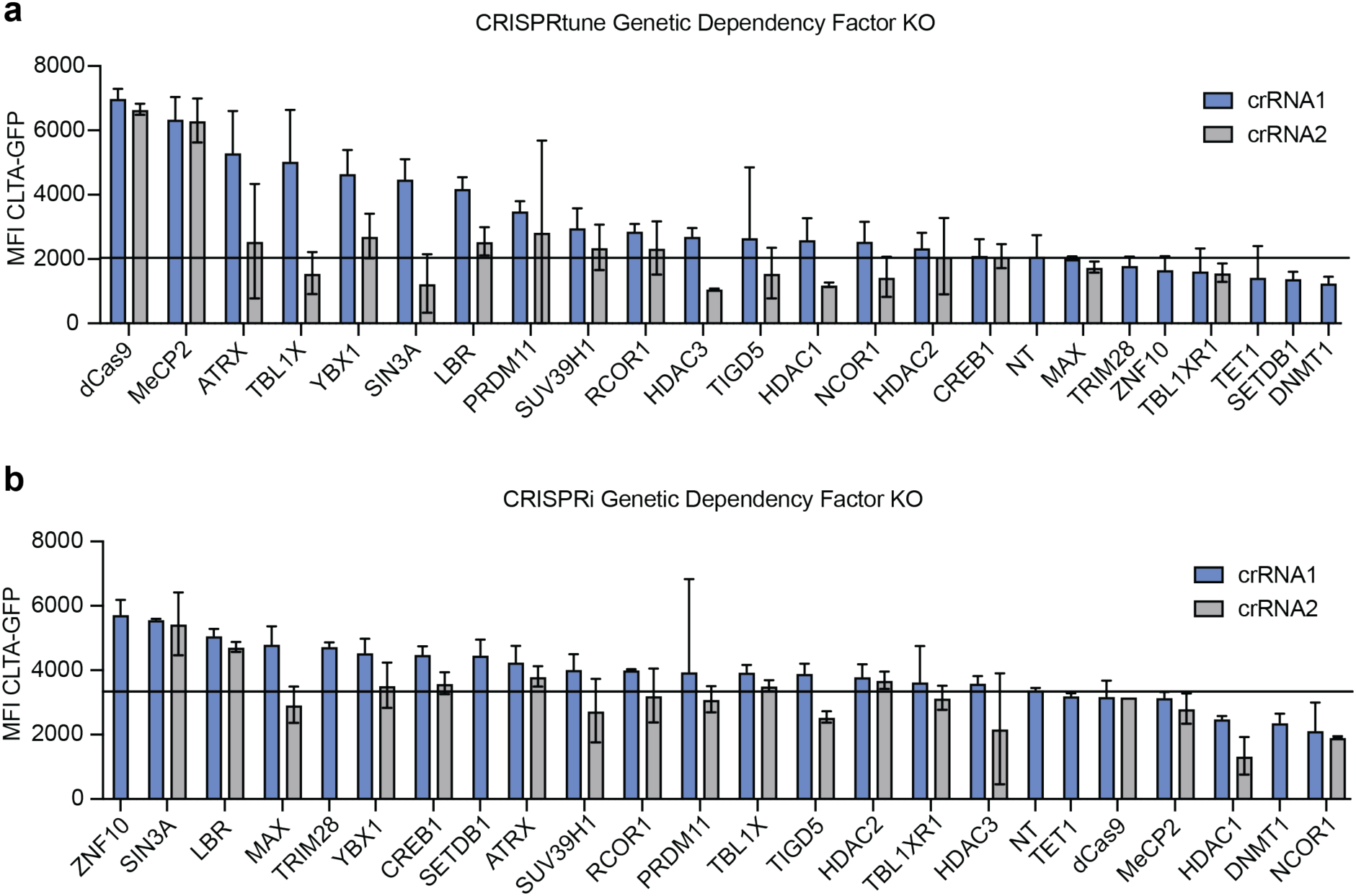
Targeted knockouts reveal MeCP2 dependency partners involved in mediating repression. **(a,b)** Quantification of CLTA-GFP median fluorescence intensity (MFI) 10 days post introduction of AsCas12a crRNAs in HEK293T cells by lentiviral transduction. Cells constitutively express CRISPRtune (**a**) and CRISPRi (**b**) that actively silences CLTA. Cells also constitutively express enAsCas12a. crRNA1 (blue) is the crRNA with the higher MFI than crRNA2 (gray). The black line represents the MFI of a non-targeting AsCas12a crRNA, representing no dependency factor knockout.

### Statistics & reproducibility

RNA-seq and CUT&RUN sequencing data were analyzed as described above. All other statistical analysis were conducted using Prism 10 for macOS (GraphPad) and Excel for MacOS (Microsoft). All data analyses were performed using two-sided unpaired t-tests for pairwise comparisons.

